# Evidence for a chromosomal inversion maintaining divergent plumage phenotypes between extensively hybridizing yellowhammers (*Emberiza citrinella*) and pine buntings (*E. leucocephalos*)

**DOI:** 10.1101/2023.06.13.544781

**Authors:** Ellen Nikelski, Alexander S. Rubtsov, Darren Irwin

## Abstract

In an allopatric speciation model, populations of a species become isolated by a geographic barrier and develop reproductive isolation through genetic differentiation. When populations meet in secondary contact, the strength of evolved reproductive barriers determines the extent of hybridization and whether the populations will continue to diverge or merge back together. The yellowhammer (*Emberiza citrinella*) and pine bunting (*E. leucocephalos*) are avian sister species that diverged in allopatry during the Pleistocene glaciations. Though they differ greatly in plumage and form distinct genetic clusters in allopatry, these taxa show negligible mitochondrial DNA differentiation and hybridize extensively in sympatry lending uncertainty to the state of reproductive isolation in the system. To assess the strength of reproductive barriers between taxa, we examined genomic differentiation across the yellowhammer and pine bunting system. We found that extensive admixture has occurred in sympatry, indicating that reproductive barriers between taxa are weak. We also identified a putative Z chromosome inversion that underlies plumage variation in the system, with the “pine bunting” inversion form showing dominance over the “yellowhammer” form. Our results suggest that yellowhammers and pine buntings are currently at a crossroads and that evolutionary forces may push this system towards either continued differentiation or population merging. However, even if these taxa merge, recombination suppression between chromosome Z inversion forms may maintain divergent plumage phenotypes within the system. In this way, our findings highlight the important role hybridization plays in increasing the genetic and phenotypic variation as well as the evolvability of a system.

## Introduction

Speciation is a complex process requiring the evolution of pre-and post-zygotic reproductive barriers that hinder hybridization and gene flow between diverging populations (Mayr, 1942; Coyne & Orr, 2004; Price, 2008). Pre-zygotic reproductive barriers manifest prior to hybrid zygote formation and prevent successful interbreeding between divergent taxa. An example of a pre-zygotic barrier is when different populations display breeding behaviour at asynchronous times such that interbreeding is reduced compared to random expectations (e.g. Moore et al. 2005; Danley et al. 2007). Post-zygotic reproductive barriers manifest following hybrid zygote formation and cause inviability, sterility or low fitness in hybrids resulting in them producing fewer offspring than non-hybrid individuals. Bateson-Dobzhansky-Muller genetic incompatibilities (Bateson, 1909; Dobzhansky, 1937; Muller, 1942) function in this respect. Here, incompatible combinations of co-evolved parental alleles inherited by hybrid individuals decrease hybrid fitness and prevent gene flow between divergent taxa. Pre-and post-zygotic reproductive barriers are highly variable in nature, involving numerous morphological, physiological, ecological and genetic traits. With time, these diverse barriers accumulate until complete reproductive isolation is achieved between taxa and speciation is deemed complete.

In an allopatric speciation model, a single large population becomes separated into several smaller populations by a geographic barrier (Mayr, 1942; Coyne & Orr, 2004; Price, 2008). In isolation, these populations diverge genetically as a result of genetic drift and natural selection acting in the absence of homogenizing gene flow. This genetic differentiation mediates divergence in various characteristics that may act as reproductive barriers between the groups.

Once the geographic barrier is removed, the diverged populations can interact within a zone of secondary contact where the net strength of any evolved reproductive barriers is evident from the degree of hybridization and gene flow between taxa (Barton & Hewitt, 1985; 1989; Hewitt, 1988; Gompert et al. 2017). Strong reproductive barriers limit hybridization and produce narrow, stable hybrid zones. In this scenario, populations will likely continue to diverge to the point that interbreeding ceases entirely. Weak reproductive barriers will allow extensive hybridization and produce wide, unstable hybrid zones. In the face of this unrestricted gene flow, populations will merge back into one large, interbreeding group. Hence hybrid zones provide a view of the intermediate steps of speciation as well as insight into the specific evolutionary forces that mediate the delicate balance between population divergence and merging.

In most cases, reproductive barriers are based on interactions between multiple genes. Interbreeding and recombination tend to disassociate the co-evolved alleles of important barrier genes, potentially leading to reproductive barrier breakdown and population merging (reviewed in Ortiz-Barrientos et al. 2016). The maintenance of reproductive barriers between groups might be directly tied to genomic structures that suppress recombination—specifically chromosomal inversions (Noor et al. 2001; Rieseberg, 2001). Because different forms of an inversion usually do not recombine, the alleles responsible for a reproductive barrier are able to accumulate within these structures and remain linked together in the face of hybridization. In this way, reproductive barriers can be retained between hybridizing populations that each possess a different orientation of the inversion. In a few study systems there is evidence for an association between inversions and the genes responsible for reproductive barriers (Lowry & Willis, 2010; Ayala et al. 2013; Todesco et al. 2020).

Colouration traits, particularly male colouration traits, are one group of characteristics that has gained special attention for their presumed ability to act as either pre-or post-zygotic reproductive barriers. Divergence in male colouration in conjunction with female preference can increase discrimination during mate choice and decrease the likelihood of interbreeding between taxa, hence acting as a pre-zygotic barrier (West-Eberhard, 1983; Grant & Grant, 1997; Edwards et al. 2005). Alternatively, hybrid individuals may be less attractive to potential mates due to intermediate or transgressive colouration patterns, resulting in a post-zygotic barrier (Bridle et al. 2006; Irwin, 2020). In line with both of these ideas, there is much evidence of divergent colouration patterns acting as reproductive barriers between taxa (e.g. Saetre et al. 1997; Seehausen & van Alphen, 1998; van der Sluijs et al. 2008; Uy et al. 2009; Turbek et al. 2021). Nevertheless, in other systems, strong colour polymorphisms are maintained within species (e.g. Tuttle, 2003; Wang & Shaffer, 2008; Hedrick et al. 2016) which lends some uncertainty to the importance of colouration traits in the speciation process.

Much recent research investigating the role of colouration traits as reproductive barriers has focused on determining the genetic underpinnings of these characteristics to better understand how specific colouration genes differentiate during speciation. This work has been particularly successful in identifying genes that regulate melanin-based traits such as melanocortin-1 receptor (MC1R; Theron et al. 2001; Rosenblum et al. 2004; Gross et al. 2009) and agouti-signaling protein (ASIP; Kingsley et al. 2009; Cerdà-Reverter et al. 2005; Haupaix et al. 2018). Carotenoid-based traits have proved more challenging to study as these pigments are obtained from the environment and deposited on the integument rather than produced endogenously like melanin (Hubbard et al. 2010; Mason & Bowie, 2020). While some genes have been implicated in carotenoid deposition (e.g., beta-carotene oxygenase 2 [BCO2; Toews et al. 2016; Andrade et al. 2019] and “scavenger receptor” genes [Brelsford et al. 2017; Toomey et al. 2017]), further work is needed to understand the genetics of colouration traits and their link to reproductive barriers between taxa.

An excellent opportunity to investigate relationships between phenotypic variation, genomic differentiation, and the processes of population divergence or merging is provided by the yellowhammer (Passeriformes: Emberizidae: *Emberiza citrinella*) and pine bunting (*E. leucocephalos*) system—a sister pair of Palearctic songbird species that show large differences in plumage colouration (Panov et al. 2003; Rubtsov and Tarasov, 2017). Hypothesized to have diverged in allopatry during the Pleistocene glaciations, these taxa currently occupy opposite sides of Eurasia and hybridize extensively within a large secondary contract zone in central and western Siberia (Panov et al. 2003; 2007; Rubtsov, 2007; Rubtsov & Tarasov, 2017; Figure 1A). Individuals within this contact zone show variable plumage phenotypes ranging from a pure yellowhammer phenotype (bright yellow body plumage and minimal facial markings) to a pure pine bunting phenotype (white body plumage and chestnut facial markings), along with various intermediate, presumed hybrid phenotypes (Figure 1B). Previous genomic work showed that yellowhammers and pine bunting have essentially undifferentiated mitochondrial DNA (mtDNA)—likely as a result of adaptive mtDNA introgression (Irwin et al. 2009)—and presented evidence for co-introgression of mitonuclear genes (Nikelski et al. 2023). A survey of nuclear genetic differentiation found several genomic regions of high differentiation that stood out against the weak differentiation seen across much of the genome (Nikelski et al. 2023). In particular, there is a wide region on chromosome Z that is highly differentiated between allopatric yellowhammers and pine buntings. These conflicting pictures of substantial plumage divergence and genetic distinctness in allopatry versus mtDNA introgression and widespread hybridization in sympatry pose the question of how strong reproductive barriers are between yellowhammers and pine buntings.

**Figure 1.**
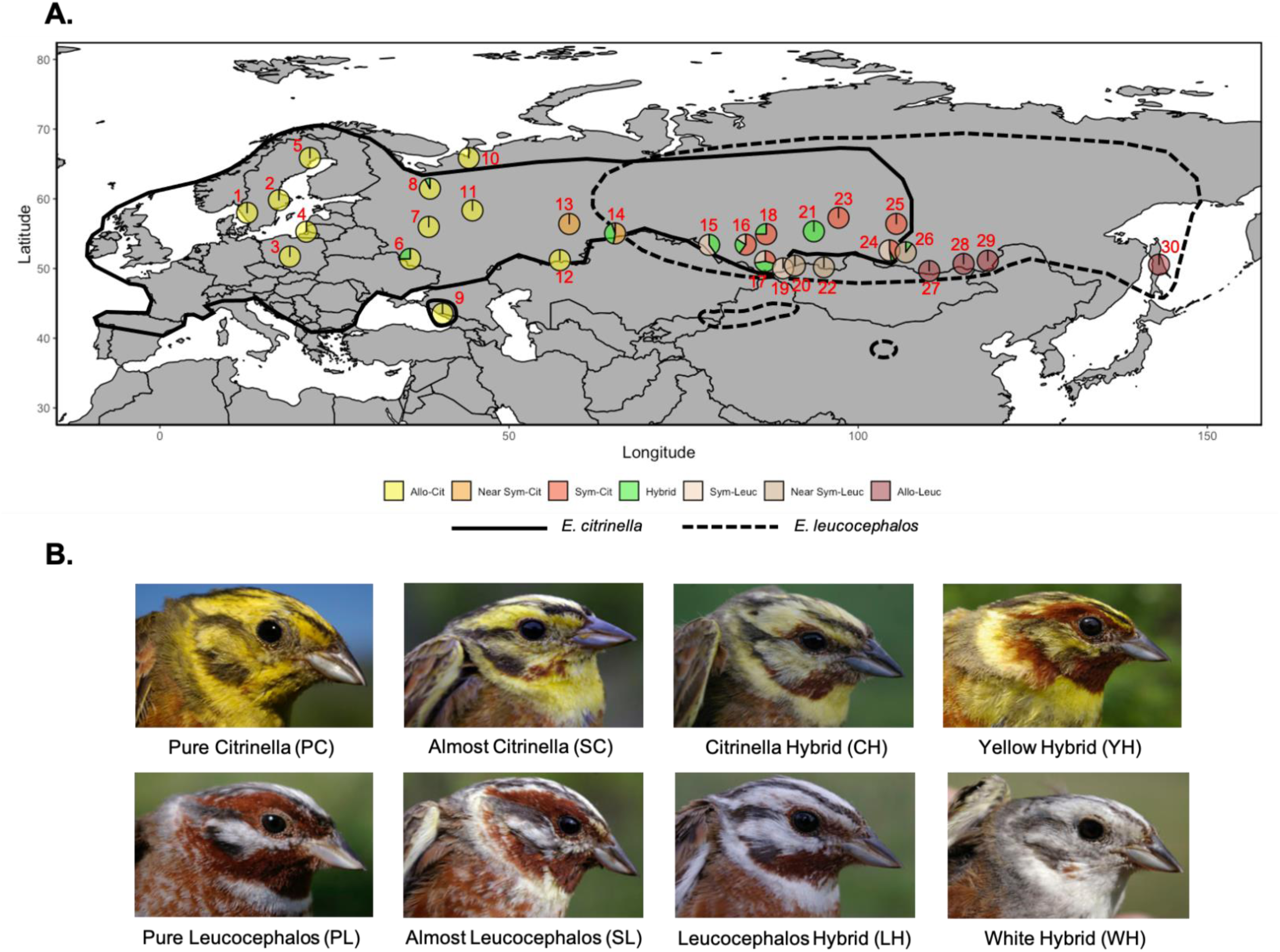
**A)** Map of sampling locations. Red numbers accompanying each sampling location pie correspond to the sampling location numbers that appear in Table 1, which shows sample sizes and sample compositions. Sampling locations may include multiple sites that appeared too close together to be shown in detail on the map. Full details for the sites included in each sampling locations can be found in Supplementary Table 1. Each sampling location pie is coloured and divided based on the proportion of each sample type that appeared within it. The sample types include: allopatric yellowhammers (Allo-Cit; yellow), near-sympatric yellowhammers (Near Sym-Cit; light orange), sympatric yellowhammers (Sym-Cit; red-orange), hybrids (Hybrid; green), sympatric pine buntings (Sym-Leuc; peach), near sympatric leucocephalos (Near Sym-Leuc; taupe) and allopatric pine buntings (Allo-Leuc; brown). The solid black line indicates the geographic range of the yellowhammer and the dashed black line indicates the geographic range of the pine bunting as described in Irwin et al. (2009). This map has been modified from Figure 1A in Nikelski et al. (2023). **B)** Photos depicting phenotypic variation within the yellowhammer and pine bunting system. Each photo represents one of eight phenotypic classes that individuals are divided into based on variation at three plumage traits: background plumage colour, amount of chestnut plumage at the brow and amount of chestnut plumage at the throat. The photos show one example of each class, but are unable to capture the full variation within each phenotypic group. Some photos appeared previously in Nikelski et al. 2023. All photos are credited to Dr. Alexander Rubtsov.

In this study, we analyzed genomic variation across the entire yellowhammer and pine bunting system to determine the genetic basis for phenotypic differentiation and the strength of potential reproductive barriers. First, we compared genomic differentiation patterns of phenotypically pure and hybrid individuals near and within the sympatric zone against what was seen between the allopatric zones. Separation of sympatric yellowhammers and pine buntings into distinct genetic clusters similar to what was seen for allopatric populations (Nikelski et al. 2023) would support the existence of strong reproductive barriers between taxa, whereas extensive genetic admixture between these sympatric populations would suggest weak reproductive barriers. To provide insight into the dynamics of hybridization and how phenotypic variation relates to genomic ancestry, we examined genetic differentiation across the different sympatric yellowhammer, pine bunting and hybrid phenotypic classes. In particular, we investigated whether the highly differentiated region identified on chromosome Z between allopatric populations (Nikelski et al. 2023) was retained within the sympatric zone or homogenized by gene flow. Maintenance of this “island of differentiation” could suggest that it houses loci important to reproductive barriers between taxa and/or that it contains a genomic structure that suppresses recombination such as a chromosomal inversion. Finally, we examined the genetic underpinnings of plumage differences between yellowhammers, pine buntings and hybrids using admixture mapping. Plumage traits are the main way by which yellowhammers and pine buntings are currently classified, but it is unknown whether this variation represents an important reproductive barrier between taxa or is merely a polymorphism of the system. Thus, we aim to use our findings regarding genomic differentiation, phenotypic variation and the state of reproductive isolation to predict the fate of yellowhammer and pine bunting system which shows potential for either populations divergence or population merging.

## Materials and Methods

### Sampling

Blood and tissue samples were collected from 321 individuals: 135 phenotypic yellowhammers (hereafter “yellowhammers”), 112 phenotypic pine buntings (“pine buntings”) and 74 phenotypic hybrids (“hybrids”; Figure 1A; Table 1; Supplementary Table 1). Genetic information from a subset of these samples was analyzed previously, for other purposes, by Irwin et al. (2009) and by Nikelski et al. (2023), whereas 148 individuals were analyzed for the first time in the present study.

**Table 1.** Geographic locations, sampling sizes and phenotypic categories for each of the sites included in this study. The sampling location umbers that appear in the “Sampling Location” column correspond to those that appear in red in Figure 1A. Sampling locations may include ultiple sites that appeared too close together to be shown in detail in Figure 1A. Full details for the sites included in each sampling location can be found in Supplementary Table 1. The “Sample Size” column describes the total number of samples collected from a particular site. Columns “Allopatric E. cit” – “Allopatric E. leuc” describe the demographic breakdown of samples within each sampling location. “E. cit.” presents *Emberiza citrinella* or yellowhammers and “E. leuc.” represents *Emberiza leucocephalos* or pine buntings.

| Sampling Location | Latitude (°N) | Longitude (°E) | Sample Size | Allopatric E. cit. | Near Sympatric E. cit. | Sympatric E. cit. | Hybrid | Sympatric E. leuc. | Near Sympatric E. leuc. | Allopatric E. leuc |
| --- | --- | --- | --- | --- | --- | --- | --- | --- | --- | --- |
| 1 | 57.99 | 12.49 | 1 | 1 | 0 | 0 | 0 | 0 | 0 | 0 |
| 2 | 59.81 | 17.05 | 1 | 1 | 0 | 0 | 0 | 0 | 0 | 0 |
| 3 | 51.71 | 18.61 | 1 | 1 | 0 | 0 | 0 | 0 | 0 | 0 |
| 4 | 55.28 | 20.97 | 5 | 5 | 0 | 0 | 0 | 0 | 0 | 0 |
| 5 | 65.86 | 21.48 | 2 | 2 | 0 | 0 | 0 | 0 | 0 | 0 |
| 6 | 51.38 | 35.84 | 4 | 3 | 0 | 0 | 1 | 0 | 0 | 0 |
| 7 | 55.97 | 38.50 | 18 | 18 | 0 | 0 | 0 | 0 | 0 | 0 |
| 8 | 61.45 | 38.67 | 13 | 12 | 0 | 0 | 1 | 0 | 0 | 0 |
| 9 | 43.54 | 40.47 | 1 | 1 | 0 | 0 | 0 | 0 | 0 | 0 |
| 10 | 65.85 | 44.24 | 1 | 1 | 0 | 0 | 0 | 0 | 0 | 0 |
| 11 | 58.33 | 44.76 | 1 | 1 | 0 | 0 | 0 | 0 | 0 | 0 |
| 12 | 51.20 | 57.27 | 7 | 7 | 0 | 0 | 0 | 0 | 0 | 0 |
| 13 | 56.39 | 58.62 | 5 | 0 | 5 | 0 | 0 | 0 | 0 | 0 |
| 14 | 55.03 | 65.19 | 19 | 0 | 10 | 0 | 8 | 0 | 1 | 0 |
| 15 | 53.39 | 78.65 | 5 | 0 | 0 | 0 | 2 | 3 | 0 | 0 |
| 16 | 53.42 | 83.89 | 25 | 0 | 0 | 15 | 6 | 4 | 0 | 0 |
| 17 | 51.04 | 86.68 | 120 | 0 | 0 | 35 | 52 | 33 | 0 | 0 |
| 18 | 54.93 | 86.82 | 4 | 0 | 0 | 3 | 1 | 0 | 0 | 0 |
| 19 | 50.02 | 89.23 | 2 | 0 | 0 | 0 | 0 | 2 | 0 | 0 |
| 20 | 50.42 | 90.93 | 3 | 0 | 0 | 0 | 0 | 0 | 3 | 0 |
| 21 | 55.33 | 93.65 | 1 | 0 | 0 | 0 | 1 | 0 | 0 | 0 |
| 22 | 50.18 | 95.09 | 5 | 0 | 0 | 0 | 0 | 0 | 5 | 0 |
| 23 | 57.28 | 97.18 | 1 | 0 | 0 | 1 | 0 | 0 | 0 | 0 |
| 24 | 52.6 | 104.48 | 19 | 0 | 0 | 8 | 1 | 10 | 0 | 0 |
| 25 | 56.41 | 105.46 | 5 | 0 | 0 | 5 | 0 | 0 | 0 | 0 |
| 26 | 52.33 | 106.81 | 10 | 0 | 0 | 0 | 1 | 0 | 9 | 0 |
| 27 | 49.64 | 110.17 | 2 | 0 | 0 | 0 | 0 | 0 | 0 | 2 |
| 28 | 50.66 | 115.09 | 17 | 0 | 0 | 0 | 0 | 0 | 0 | 17 |
| 29 | 51.12 | 118.56 | 15 | 0 | 0 | 0 | 0 | 0 | 0 | 15 |
| 30 | 50.56 | 143.08 | 8 | 0 | 0 | 0 | 0 | 0 | 0 | 8 |
| **Total** |  |  | **321** | **53** | **15** | **67** | **74** | **52** | **18** | **42** |

When possible, sampled birds were photographed and males were scored based on protocols presented in Panov et al. (2003) and Rubtsov and Tarasov (2017) for three morphological traits that distinguish yellowhammer and pine bunting phenotypes: background plumage colour, amount of chestnut at the brow (vs. yellow or white) and amount of chestnut at the throat (vs. yellow or white). Background colour ranged from bright yellow to pure white and was assessed for head and body regions that did not show brown or black streaking. Males were given a score from 0-7 for each trait with scores of 0 indicating a pure yellowhammer phenotype and scores of 7 indicating a pure pine bunting phenotype. Following scoring, males were sorted into one of eight phenotypic classes—pure *citrinella* (PC), almost *citrinella* (SC, for “semi-*citrinella*”), *citrinella* hybrid (CH), yellow hybrid (YH), white hybrid (WH), *leucocephalos* hybrid (LH), almost *leucocephalos* (SL, for “semi-*leucocephalos*”) and pure *leucocephalos* (PL; Figure 1B). Unless otherwise indicated, PC and SC individuals were grouped as phenotypic yellowhammers, SL and PL individuals were grouped as phenotypic pine buntings, and the remaining classes were categorized as hybrids in analyses (Rubtsov and Tarasov, 2017). Female individuals do not show the same plumage variation as males and therefore were not phenotypically scored or included in analyses of phenotypic variation. Yellowhammer, pine bunting and hybrid individuals were further classified by geographic location. “Sympatric” individuals were sampled within the sympatric zone (Figure 1A). “Near-sympatric” individuals were sampled outside of but close to the border of the sympatric zone such that a recent expansion of this area might capture these regions. The possibility of an expansion is supported by the dynamic nature of the sympatric zone (Panov et al. 2003; 2007; Rubtsov, 2007; Irwin et al. 2009; Rubtsov & Tarasov, 2017) and by a small number of hybrids found within the near-sympatric zone. “Allopatric” individuals were sampled far (more than 400 kilometres) from the border of the sympatric zone.

### DNA sequencing and identification of single nucleotide polymorphisms DNA extraction and genotyping-by-sequencing

We used a standard phenol-chloroform method (Sambrook et al. 1989) to extract DNA from blood and tissue samples. DNA was resuspended in 1× TE buffer and stored at 4°C until library preparation. We prepared four genotyping-by-sequencing (GBS) libraries (Elshire et al. 2011) that together included DNA from the 321 samples discussed above and an additional 14 samples from other members of Emberizidae that were analyzed in Nikelski et al. (2023). Library preparation followed the procedures described in Nikelski et al. (2023).

### Genotyping-by-sequencing data filtering

We conducted GBS read filtering as per the protocols described in Nikelski et al. (2023). Briefly, reads were demultiplexed, trimmed and aligned to the zebra finch reference genome (*Taeniopygia guttata* version 3.2.4; Warren et al. 2010). Afterwards, single nucleotide polymorphisms (SNPs) were identified between individuals, resulting in a genome-wide VCF file. The loci contained within this file were filtered for quality. Depth of coverage averaged 16.11 reads per SNP per individual.

### Variant site analyses

We employed Admixture (Alexander et al. 2009) version 1.3.0 to estimate the number of ancestral populations in the yellowhammer and pine bunting system and to assign ancestry proportions to each sample. Before running this program, a VCF file of SNP information from only yellowhammers, pine buntings and hybrids was created and trimmed for linkage disequilibrium using Plink version 1.9 (Chang et al. 2015), isolating 417,164 SNPs for analysis. Within Admixture, we ran six maximum likelihood models with “K” values ranging from 1 to 6 clusters. A run was terminated when the difference between the log-likelihood values of two consecutive iterations dropped below 1 × 10^−10^. Of the six models, cross-validation error was lowest for “K=1”, but was similarly low for “K=2”. Based on the phenotypic, behavioural and ecological differences seen between yellowhammers and pine buntings (Panov et al. 2003; Rubtsov & Tarasov, 2017) as well as the genetic differentiation identified between allopatric populations (Nikelski et al. 2023), we plotted the ancestry proportions produced for “K=2”.

We loaded the genome-wide VCF file of all Emberizidae individuals into R (R Core Team, 2014) version 3.6.2. Versions of the scripts presented in Irwin et al. (2018) were used to filter variant sites and to calculate the sample size, allele frequency and Weir and Cockerham’s *F*_ST_ (Weir and Cockerham, 1984) for each SNP when comparing allopatric yellowhammers (n = 53) and allopatric pine buntings (n = 42). A total of 374,780 SNPs passed quality thresholds and were variable among these allopatric individuals. Using this SNP information, we conducted a principal components analysis (PCA) of yellowhammers, pine buntings and hybrids using the pcaMethods package (Stacklies et al. 2007) with the “svdImpute” command to impute missing genomic data. To investigate how differentiation varied between chromosomes, we also conducted chromosome-specific PCAs.

We identified several outliers in our genome-wide PCA suggesting potential kinship between these individuals. To investigate this possibility, we used the SNPRelate package (Zheng et al. 2012) to convert a variant site VCF file of yellowhammer, pine bunting and hybrid genetic information into GDS format and calculate identity-by-descent coefficients between individuals using the KING method (Manichaikul et al. 2010). The resulting coefficients were visualized as a heatmap using the lattice package (Sarkar, 2008).

To investigate patterns of genetic variation across sympatric phenotypic classes, we extracted the PC1 value associated with each sympatric individual from the genome-wide PCA discussed above and that is shown in different forms in Figure 2A and Supplementary Figure 3. We removed sympatric females and individuals with unknown phenotypes and then plotted the PC1 values as a series of boxplots based on phenotypic class. Similarly, we also extracted the ancestry proportions produced with Admixture using “K=2” for the same set of sympatric individuals and plotted the proportions based on the phenotypic class.

**Figure 2.**
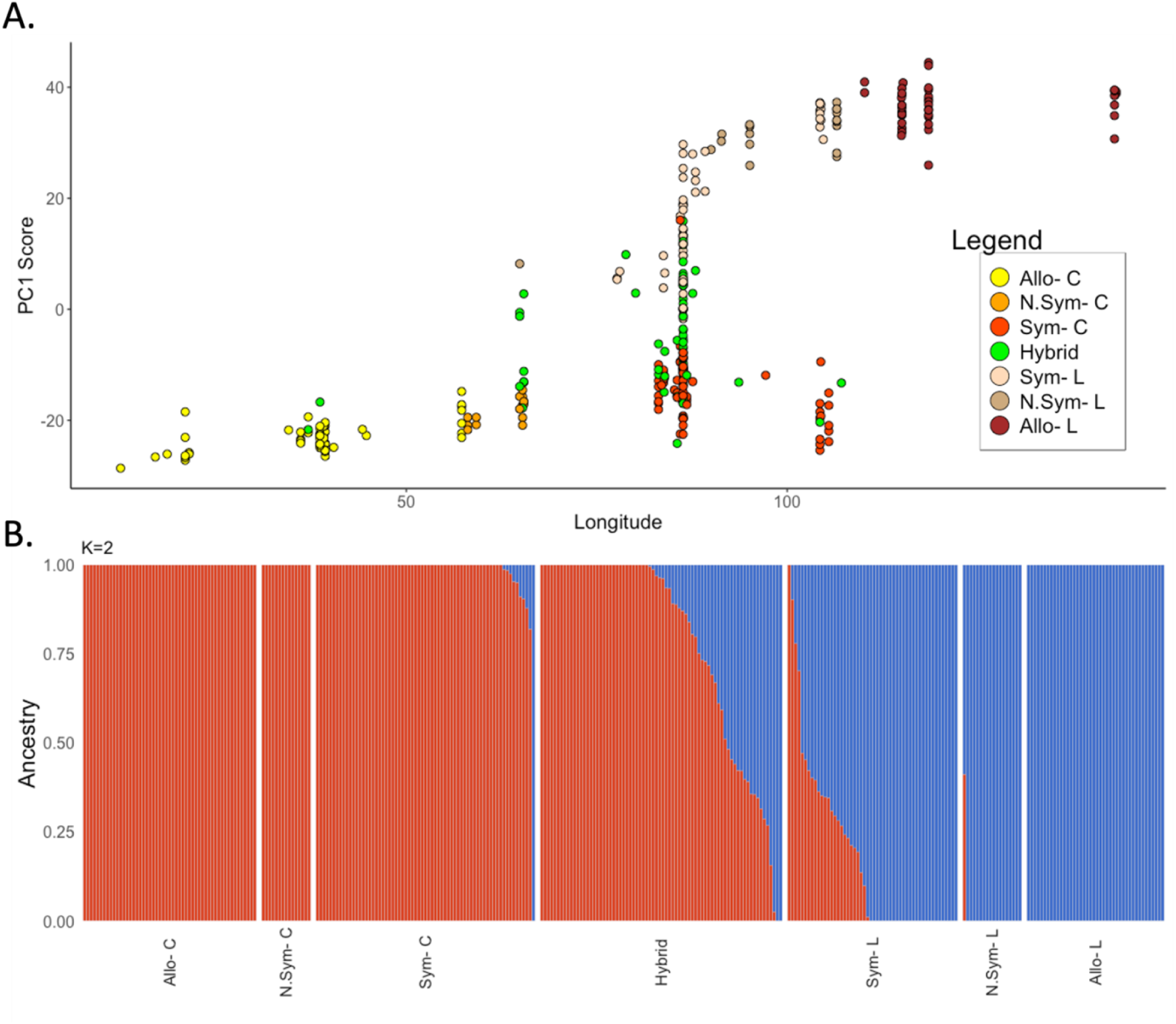
Genetic relationships between allopatric yellowhammers (Allo-C, n = 53), near-sympatric yellowhammers (N.Sym-C, n = 15), sympatric yellowhammers (Sym-C, n =67), allopatric pine buntings (Allo-L, n = 42), near-sympatric pine buntings (N.Sym-L, n = 18), sympatric pine buntings (Sym-L, n = 52) and hybrids (n = 74). **A)** PC1 values for allopatric yellowhammers (yellow), near-sympatric yellowhammers (light orange), sympatric yellowhammers (red-orange), allopatric pine buntings (brown), near-sympatric pine buntings (taupe), sympatric pine buntings (peach) and hybrids (green) from a PCA comparing information from 374,780 SNPs between allopatric yellowhammers and pine buntings (Supplementary Figure 3) plotted against the longitude at which each individual was sampled. PC1 explained explains 1.4% if the variation among individuals. **B)** Ancestry proportions of allopatric yellowhammers, near-sympatric yellowhammers, sympatric yellowhammers, allopatric pine buntings, near-sympatric pine buntings, sympatric pine buntings and hybrids as predicted by an Admixture model with K=2. Information from 417,164 SNPs were included in this analysis.

### Chromosome Z analyses

Based on unanticipatedly strong population structure identified in a chromosome Z PCA when comparing all samples, we extracted a subset of individuals from our dataset to be graphed in “genotype-by-individual” plots. The code for this plot is included in scripts provided by Irwin et al. (2018) although such a figure did not appear in that publication.

Results from genotype-by-individual graphs suggested tight linkage between high *F_ST_* loci in a particular section of chromosome Z. To investigate this region for a potential chromosomal inversion, we conducted a linkage disequilibrium analysis of this chromosome. First, we used VCFtools (Danecek et al. 2011) to create three VCF files of chromosome Z SNP information from male individuals that were separated based on the groups shown in Z chromosome genotypic clusters. Linkage disequilibrium statistics were calculated for each of the VCF files as squared inter-variant allele correlations using Plink version 1.9 (Chang et al. 2015) and the –r2 flag. Correlations were visualized in R using the ggpubr package (Kassambara, 2017).

### Plumage trait admixture mapping

We conducted admixture mapping on three plumage traits that differ between yellowhammer and pine buntings: the background colour, the amount of chestnut at the brow, and the amount of chestnut at the throat. This was completed using the program GEMMA (Zhou and Stephens, 2012) which applies a genome-wide efficient mixed model association algorithm to test associations between variation in the genome and variation in phenotypic traits. GEMMA requires genomic data to be imputed or “complete” before it is run through admixture mapping. To meet this criterion, we used the program Bimbam (Scheet and Stephens, 2006) with the parameters 10 EM, 40 steps, and 7 clusters to impute missing genotypes using genotypic information at SNPs in high linkage with the missing SNP. It should be noted that Bimbam only considers genetic linkage disequilibrium information among the cohort of samples provided. It does not take into account the different phenotypes of each individual when filling in missing genetic data. Using this imputed dataset, we created a centred relatedness matrix in GEMMA to be used in later admixture mapping. We then used GEMMA to run three univariate linear mixed models (ULMM)—one for each phenotypic trait—and used a likelihood ratio test to assess association between genomic and phenotypic information at each SNP. Based on GEMMA’s internal filtering parameters, 220,220 SNPs were included in the background ULMM, 220,124 SNPs were included in the brow ULMM and 220,307 SNPs were included in the throat ULMM. Only male individuals with phenotypic scores at all traits (n = 230) were included in admixture mapping. Probability values from the likelihood ratio tests performed in each ULMM were visualized on Manhattan plots created with the “qqman” package in R (Turner, 2018).

### Investigation of plumage trait dominance

Following admixture mapping, we investigated patterns of dominance for each of the SNPs significantly associated with a plumage trait of interest. In particular, we focused on SNPs located within a putative chromosomal inversion located on chromosome Z that were strongly associated with facial plumage variation and weakly associated with body plumage variation. To conduct this analysis, we created contingency tables that compared the phenotypic scores of each trait against the genotypes of significantly associated SNPs. These tables were visualized as balloon plots using the ggpubr package (Kassambara, 2017) and examined for patterns suggestive of dominance.

## Results

### Genetic variation inside and outside the hybrid zone

Our analysis showed yellowhammers and pine buntings sampled far from the hybrid zone as being genetically distinct, consistent with the results of Nikelski et al. (2023) and near-sympatric populations as being slightly less genetically differentiated from each other (Figure 2A; Supplementary Figure 1).

Moving into the sympatric zone, we saw a partial breakdown in the genetic distinctiveness of phenotypic yellowhammers and pine buntings (Figure 2; Supplementary Figure 2). The program Admixture estimated that, while the majority of phenotypic yellowhammers showed putatively pure yellowhammer genetic ancestry, ten individuals showed some pine bunting ancestry (Figure 2B). Phenotypic pine buntings were estimated to show even greater admixture within the sympatric zone with roughly half of the individuals possessing anywhere between 1-78% putative yellowhammer genetic ancestry and the other half possessing putatively pure pine bunting genetic ancestry (Figure 2B). Two phenotypic pine buntings were estimated to possess more than 90% yellowhammer ancestry despite their pure plumage type.

Phenotypic hybrids were also varied in their genotypes. Many phenotypic hybrids displayed intermediate ancestries with genetic input from both yellowhammer and pine bunting populations (Figure 2). Nevertheless, a large portion of the phenotypic hybrids possessed greater yellowhammer genetic ancestry and tended to group more closely with phenotypic yellowhammers than with phenotypic pine buntings in a PCA (Figure 2A)—a pattern that was reported previously in this system (Irwin et al. 2009). A total of 33 phenotypic hybrids were estimated as having putatively pure yellowhammer ancestry despite their intermediate plumage patterns while only two showed putatively pure pine bunting ancestry (Figure 2B). Very few individuals showed genetic ancestry proportions close to 50-50 as might be expected in F1 hybrids.

Grouping sympatric individuals based on their phenotypic classes provided further insight into genetic variation and hybridization within the sympatric zone. The eight phenotypic classes showed a continuous distribution of PC1 values (Figure 3A) and Admixture estimated ancestry scores (Figure 3B), with a strong association between genetic score and phenotypic class. PC1 values from “PC” and “PL” individuals appeared at the extremes of this distribution while “SC” and “SL” individuals grouped closely with these groups, but possessed slightly more intermediate PC1 values on average. “CH” and “LH” individuals showed increasingly intermediate PC1 values followed by “YH” and “WH” individuals which formed the centre of the distribution. Yellowhammer and pine bunting genetic ancestry was represented in all eight sympatric phenotypic classes (Figure 3B). One “PC” and one “SC” individual showed some amount of pine bunting ancestry while the rest of the individuals in these groups showed putatively pure yellowhammer ancestry. For the “CH”, “YH” and “WH” phenotypic classes, most individuals showed greater yellowhammer than pine bunting genetic ancestry with several individuals in each phenotypic class showing putatively pure yellowhammer ancestry and only one “CH” individual showing putatively pure pine bunting ancestry. The majority of “LH” and “SL” individuals showed greater pine bunting ancestry with one and four individuals possessing putatively pure pine bunting ancestry respectively. Interestingly, one “SL” individual also showed putatively pure yellowhammer ancestry. Finally, over half of the “PL” individuals possessed putatively pure pine bunting genetic ancestry while the remaining individuals showed varying amounts of yellowhammer ancestry (up to 78%).

**Figure 3.**
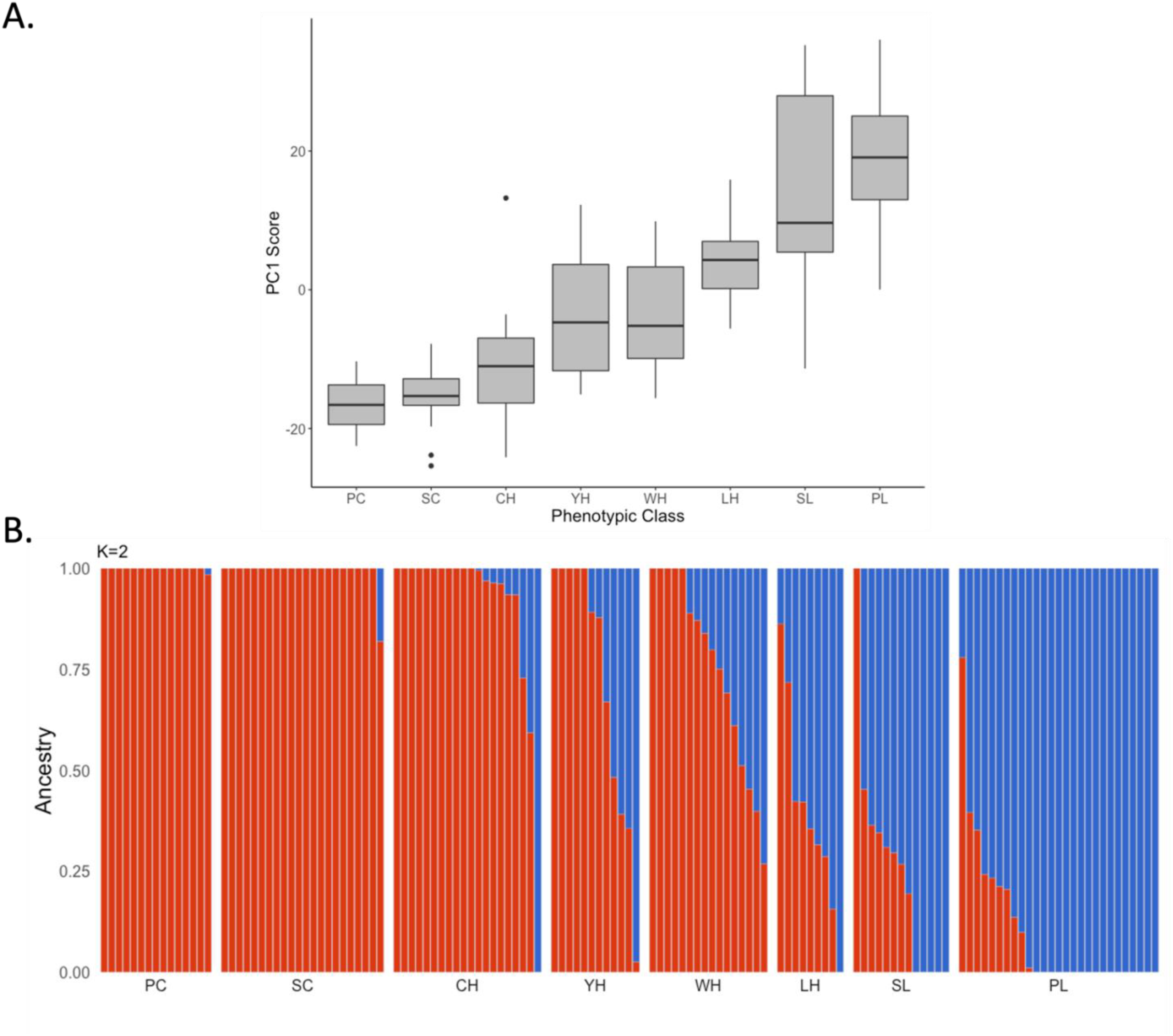
Genetic relationships between sympatric individuals of different phenotypic classes including “pure citrinella” (PC, n = 15), “almost citrinella” (SC, n = 22), “citrinella hybrid (CH, n = 20), “yellow hybrid” (YH, n = 12), “white hybrid” (WH, n = 16), almost leucocephalos (SL, n = SL) and “pure leucocephalos” (PL, n = 27) individuals. Descriptions of how individuals were sorted into these phenotypic classes are available in Panov et al. (2003) and Rubtsov and Tarasov (2017) and images of individuals within each phenotypic class are available in Figure 1B. **A)** The PC1 values of sympatric individuals extracted from a PCA comparing information from 374,780 SNPs between allopatric yellowhammers and pine buntings (Supplementary Figure 3) plotted against the phenotypic class of individuals. PC1 values for each phenotypic class are summarized as box plots. **B)** Ancestry proportions of sympatric individuals belonging to different phenotypic classes as predicted by an Admixture model with K=2. Information from 417,164 SNPs were included in this analysis.

While conducting our analysis of genetic variation across the yellowhammer and pine bunting system, we identified several outliers within our PCAs (Supplementary Figures 1-3). These outliers do not affect the overall genetic trends observed among individuals and are further examined (Supplementary Figures 4, 5; Supplementary Table 2) and discussed in the supplementary materials.

### Differentiation on the Z chromosome

In our past examination of differentiation between allopatric populations (Nikelski et al. 2023), a large region of differentiation was identified between allopatric yellowhammers and pine buntings on chromosome Z. To investigate how this genomic region may be affected by gene flow across the sympatric zone, we conducted a PCA of Z chromosome SNPs. This analysis produced six discrete clusters—three clusters along PC1 that each separated into two further clusters along PC2 (Figure 4A).

**Figure 4.**
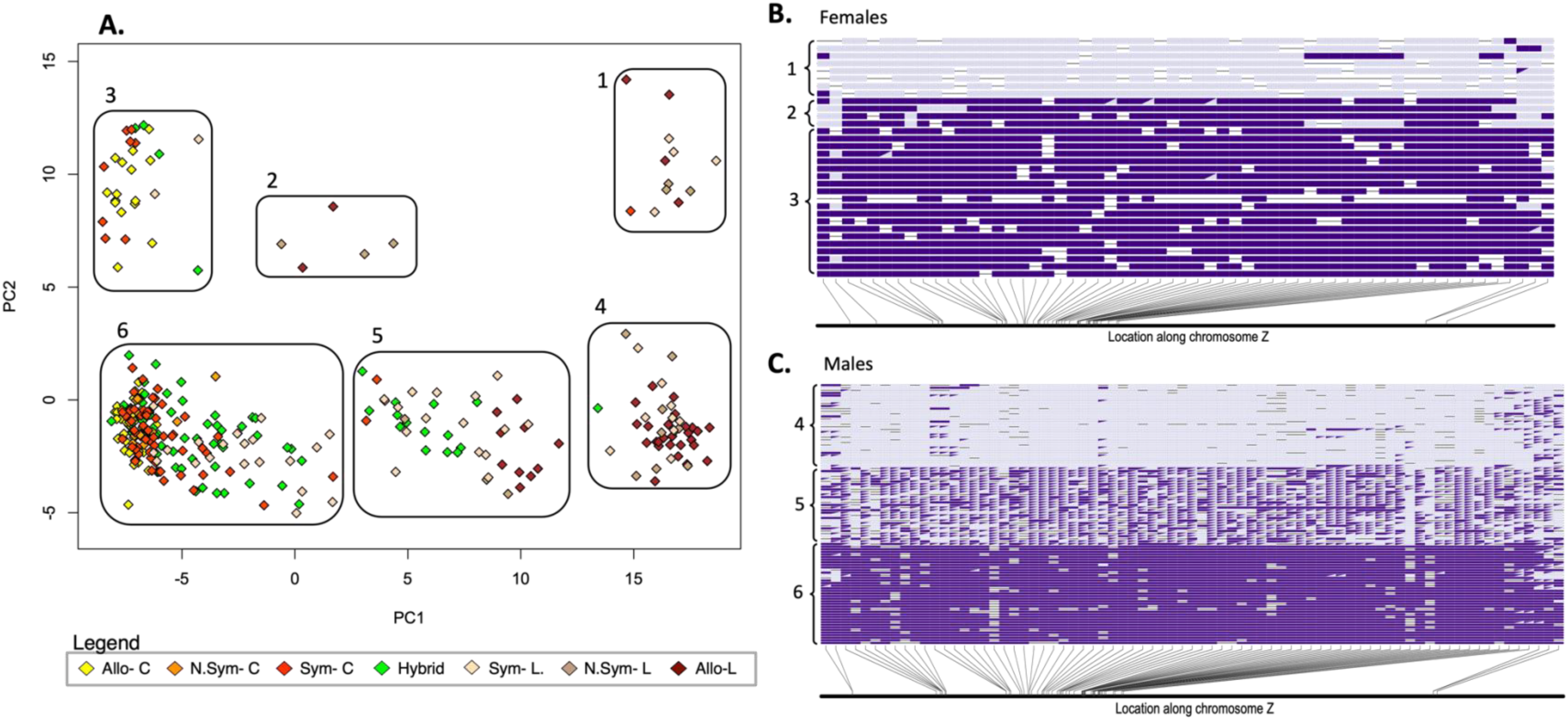
Genetic differentiation across the Z chromosome among allopatric yellowhammers (Allo-C, n = 53), near-sympatric yellowhammers (N.Sym-C, n = 15), sympatric yellowhammers (Sym-C, n = 67), allopatric pine buntings (Allo-L, n = 42), near-sympatric pine buntings (N.Sym-L, n = 18), sympatric pine buntings (Sym-L, n = 52) and hybrids (n = 74). **A)** Z chromosome PCA of allopatric yellowhammers (yellow), near-sympatric yellowhammers (light orange), sympatric yellowhammers (red-orange), allopatric pine buntings (brown), near-sympatric pine buntings (taupe), sympatric pine buntings (peach) and hybrids (green). PC1 explains 8.1% of the variation among individuals and PC2 explains 1.7% of the variation among individuals. Information from 11,147 SNPs was included in this analysis. Numbered boxes are used to designate each of 6 clusters within PC space. **B)** Genotype-by individual plot of a subset of females (n = 32) **C)** Genotype-by individuals plot of a subset of males (n = 100) from the chromosome Z PCA. SNPs with *F*ST greater or equal to 0.7 in comparisons of allopatric populations were included in this analysis. Boxes filled in with one colour indicate homozygosity at a locus and boxes split into different coloured triangles indicate heterozygosity. Light purple indicates alleles with putative pine bunting ancestry and dark purple indicates alleles with putative yellowhammer ancestry. Numbers along the left side correlate to numbered clusters within the chromosome Z PCA.

Clusters along PC1 were coarsely based on species designations, with most allopatric and near-sympatric individuals forming groups on either end of the axis, whereas sympatric and hybrid individuals tended to occur in the middle or leftmost cluster (Figure 4A). There was a fair amount of noise in this pattern. Along PC2, individuals putatively separated based on sex with all female individuals appearing in the upper three clusters and the majority of male individuals appearing in the lower three clusters (Figure 4A).

To examine the genetic drivers of the six clusters, we plotted the genotypes of high *F*_ST_ Z chromosome SNPs for a subset of individuals (Figure 4B-C). For “female” clusters (1-3), we found that cluster “1” individuals possessed pine bunting alleles at most sites while cluster “2” and “3” individuals possessed mostly yellowhammer alleles (Figure 4B). Cluster “2” and “3” individuals could be further separated based on two regions on either end of chromosome Z where cluster “2” individuals possessed pine bunting alleles and cluster “3” individuals possessed yellowhammer alleles. This latter pattern is likely responsible for the intermediate nature of cluster “2” along PC1 (Figure 4A).

For the “male” clusters (4-6), the genotypic patterns of high *F*_ST_ SNPs were more complex because they involve heterozygosity (Figure 4C). Cluster “4” individuals were homozygous for pine bunting alleles at most genomic sites while cluster “6” individuals were homozygous for yellowhammer alleles. In cluster “5” individuals, we found a large block of heterozygosity towards the middle of chromosome Z bordered by two genomic regions that were mostly homozygous for yellowhammer alleles.

Following the above analysis, we performed a linkage disequilibrium analysis of Z chromosome SNPs to confirm the presence of highly linked haploblocks within the system. When analyzing only cluster “4” individuals or only cluster “6” individuals as shown in Figure 4A, we did not find evidence of high linkage disequilibrium between distant locations on the chromosome (Figure 5A, 5C). Yet, when we analyzed individuals from all of clusters “4-6”, we identified an area of high linkage within the previously identified block of heterozygosity (Figure 5B). These results provide support for a chromosomal inversion (reviewed in Kirkpatrick, 2010) that is reducing recombination between two distinct haplotypes over a large region of chromosome Z (Nachman, 2002).

**Figure 5.**
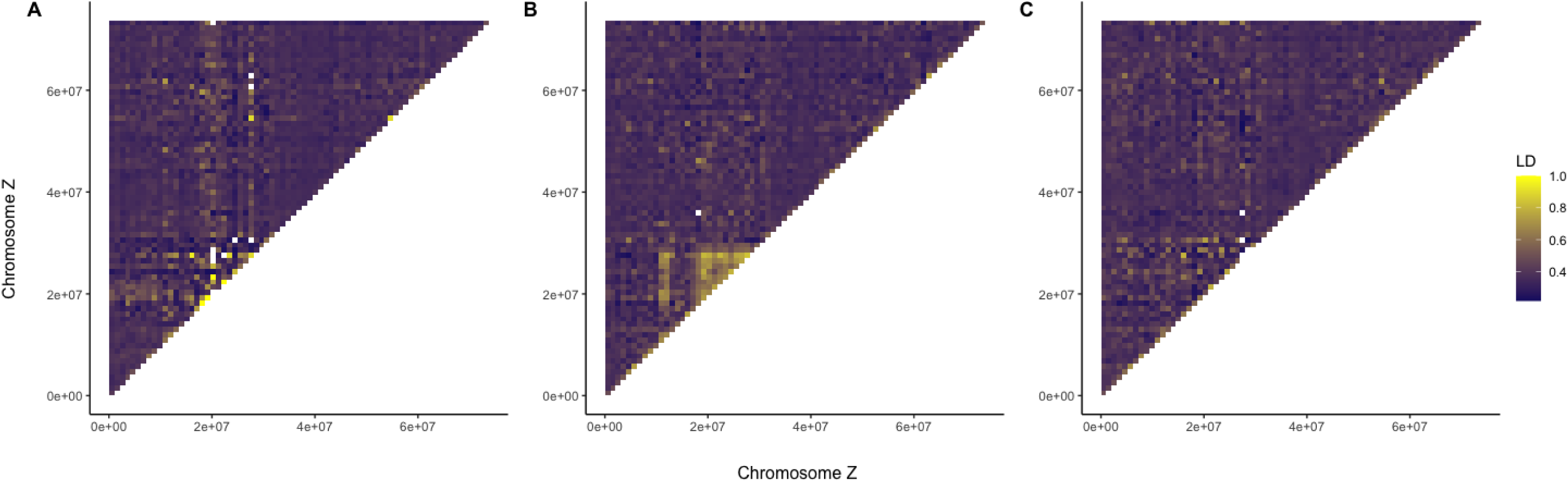
Heatmap of linkage disequilibrium across the Z chromosome based on genetic information from male individuals in **A)** Group 4, **B)** Group 4, 5 and 6, and **C)** Group 6 as classified in Figure 4A. Linkage disequilibrium statistics were calculated as squared inter-variant allele correlations between SNPs. Each tile summarizes correlations between multiple SNPs. Yellow indicates high linkage disequilibrium between sites and indigo indicates low linkage disequilibrium between sites.

### Genetic underpinnings of plumage traits

Admixture mapping revealed strong associations between specific genomic regions and three plumage traits that distinguish yellowhammers and pine buntings. Variation in body plumage colour was associated with genetic variation on chromosome 20 and chromosome Z (Figure 6A; Supplementary Figure 6A). One SNP on chromosome 20 and four SNPs on chromosome Z were significantly associated with this trait following a Bonferroni correction (Table 2), with many others showing high but nonsignificant associations. One of the significant SNPs on chromosome Z is within an annotated gene—ABCA1—and the significant SNP on chromosome 20 is within the gene CHD6.

**Figure 6.**
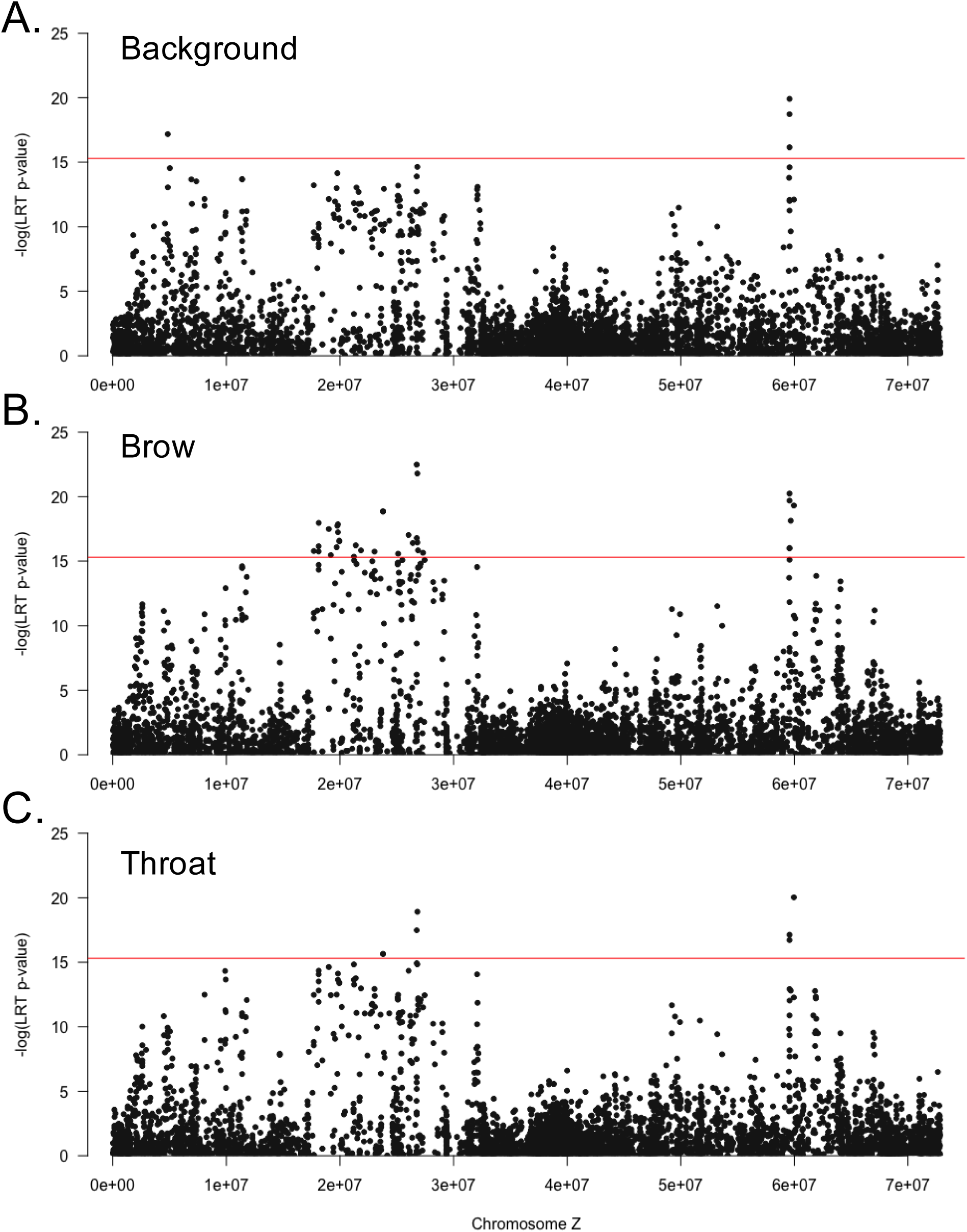
Associations between SNPs located on chromosome Z and phenotypic variation in three plumage traits within the yellowhammer and pine bunting system. P-values for each SNP were determined using a likelihood ratio test calculated in the GEMMA program. Red lines indicate Bonferroni corrected significance thresholds. **A)** Associations between 6502 chromosome Z SNPs and variation in the background plumage colour. **B)** Associations between 6509 chromosome Z SNPs and variation in the amount of chestnut plumage at the brow. **C)** Associations between 6487 chromosome Z SNPs and variation in the amount of chestnut plumage at the throat

**Table 2.**
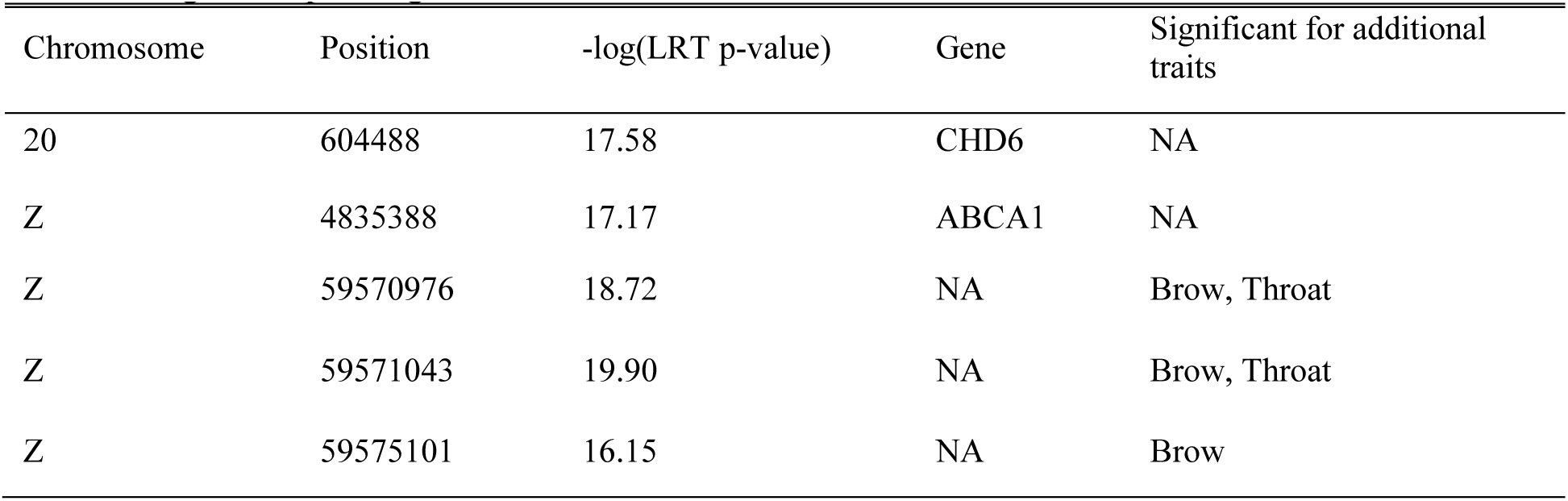
Identities of SNPs significantly associated with phenotypic variation in the background colour trait within the yellowhammer and pine bunting system. SNP locations are indicated by the “Chromosome” column which indicates the chromosomal location and the “Position” column which indicate the base pair position. P-values were calculated using a likelihood ratio test in the GEMMA program and are written in the form -log(p-value). Larger values indicate greater significance. The Bonferroni corrected significance threshold was set at 15.29811. SNPs that occur within a gene are indicated in the “Gene” column with gene names written as they appear within the zebra finch reference genome (Taeniopygia guttata version 3.2.4; Warren et al. 2010). “NA” indicates that a SNP was not found within an annotated gene. SNPs that were significantly associated with another plumage trait are indicated in the “Significant for additional traits” column where “Background” indicates the colour of the background plumage, “Brow” indicates the amount of chestnut plumage at the brow and “Throat” indicates the amount of chestnut plumage at the throat. “NA” in this column indicates that a particular significant SNP was unique to the background plumage trait.

| Chromosome | Position | $-\log(\text{LRT } p\text{-value})$ | Gene | Significant for additional traits |
| --- | --- | --- | --- | --- |
| 20 | 604488 | 17.58 | CHD6 | NA |
| Z | 4835388 | 17.17 | ABCA1 | NA |
| Z | 59570976 | 18.72 | NA | Brow, Throat |
| Z | 59571043 | 19.90 | NA | Brow, Throat |
| Z | 59575101 | 16.15 | NA | Brow |

Chestnut brow plumage showed a strong association with genetic variation on chromosome Z (Figure 6B; Supplementary Figure 6B), with 34 SNPs split between two genomic regions showing significant associations (Table 3). These include two significant SNPs within the gene GRAMD3, three within the FER gene and two within the PJA2 gene. Further annotated genes that contained a single significant SNP included: EFNA5, APC, MCC, CEP120 and ECPAS.

**Table 3.** Identities of SNPs significantly associated with phenotypic variation in the amount of chestnut plumage at the brow within the yellowhammer and pine bunting system. The Bonferroni corrected significance threshold was set at 15.29768. Additional information regarding the meaning of column headings and values can be found in the caption of Table 2.

| Chromosome | Position | -log(LRT p-value) | Gene | Significant for another phenotype? |
| --- | --- | --- | --- | --- |
| Z | 17694478 | 15.80 | NA | NA |
| Z | 18127032 | 15.75 | NA | NA |
| Z | 18127137 | 16.16 | NA | NA |
| Z | 18131016 | 17.97 | NA | NA |
| Z | 19020732 | 17.49 | NA | NA |
| Z | 19196302 | 15.47 | EFNA5 | NA |
| Z | 19708803 | 16.08 | NA | NA |
| Z | 19768210 | 17.75 | FER | NA |
| Z | 19821523 | 17.86 | FER | NA |
| Z | 19821527 | 17.24 | FER | NA |
| Z | 19939726 | 16.58 | PJA2 | NA |
| Z | 19939881 | 16.56 | PJA2 | NA |
| Z | 21223538 | 15.35 | APC | NA |
| Z | 21386283 | 16.23 | MCC | NA |
| Z | 21843152 | 15.84 | NA | NA |
| Z | 23035555 | 15.75 | CEP120 | NA |
| Z | 23774941 | 18.85 | NA | Throat |
| Z | 23774945 | 18.85 | NA | Throat |
| Z | 23774949 | 18.85 | NA | Throat |
| Z | 25116054 | 15.58 | NA | NA |
| Z | 26023933 | 17.01 | ECPAS | NA |
| Z | 26402437 | 16.41 | NA | NA |
| Z | 26757019 | 22.48 | NA | Throat |
| Z | 26760575 | 16.78 | NA | NA |
| Z | 26812200 | 16.45 | GRAMD3 | NA |
| Z | 26812248 | 21.80 | GRAMD3 | Throat |
| Z | 26869834 | 15.84 | NA | NA |
| Z | 27304119 | 15.66 | NA | NA |

| Chromosome | Position | $-\log(\text{LRT p-value})$ | Gene | Significant for another phenotype? |
| --- | --- | --- | --- | --- |
| Z | 59570976 | 19.69 | NA | Background, Throat |
| Z | 59571035 | 15.99 | NA | NA |
| Z | 59571043 | 20.24 | NA | Background, Throat |
| Z | 59575101 | 16.03 | NA | Background |
| Z | 59675542 | 18.13 | NA | NA |
| Z | 59953541 | 19.32 | NA | Throat |

Chestnut throat plumage was also strongly associated with genetic variation on the Z chromosome (Figure 6C; Supplementary Figure 6C), with eight significant SNPs (Table 4). Only one of these SNPs was found within a gene: GRAMD3.

**Table 4.** Identities of SNPs significantly associated with phenotypic variation in the amount of chestnut plumage at the throat within the yellowhammer and pine bunting system. The Bonferroni corrected significance threshold was set at 15.29851. Additional information regarding the meaning of column headings and values can be found in the caption of Table 2.

| Chromosome | Position | -log(LRT p-value) | Gene | Significant for another phenotype? |
| --- | --- | --- | --- | --- |
| Z | 23774941 | 15.64 | NA | Brow |
| Z | 23774945 | 15.64 | NA | Brow |
| Z | 23774949 | 15.64 | NA | Brow |
| Z | 26757019 | 17.47 | NA | Brow |
| Z | 26812248 | 18.91 | GRAMD3 | Brow |
| Z | 59570976 | 17.12 | NA | Background, Brow |
| Z | 59571043 | 16.72 | NA | Background, Brow |
| Z | 59953541 | 20.04 | NA | Brow |

We also investigated dominance interactions between plumage phenotypes and their related genotypes. No dominance interactions were detected for significant SNPs located on chromosome 20 and, as such, we focused our analysis on SNPs located on chromosome Z. Significant SNPs on chromosome Z possessed differing dominance patterns depending on whether they were located inside or outside of the putative chromosomal inversion discussed earlier. These patterns can be summarized by the dominance patterns seen for four SNPs highlighted in Figure 7A—two within the putative inversion, two outside the putative inversion and all highly differentiated between allopatric yellowhammers and pine buntings.

**Figure 7.**
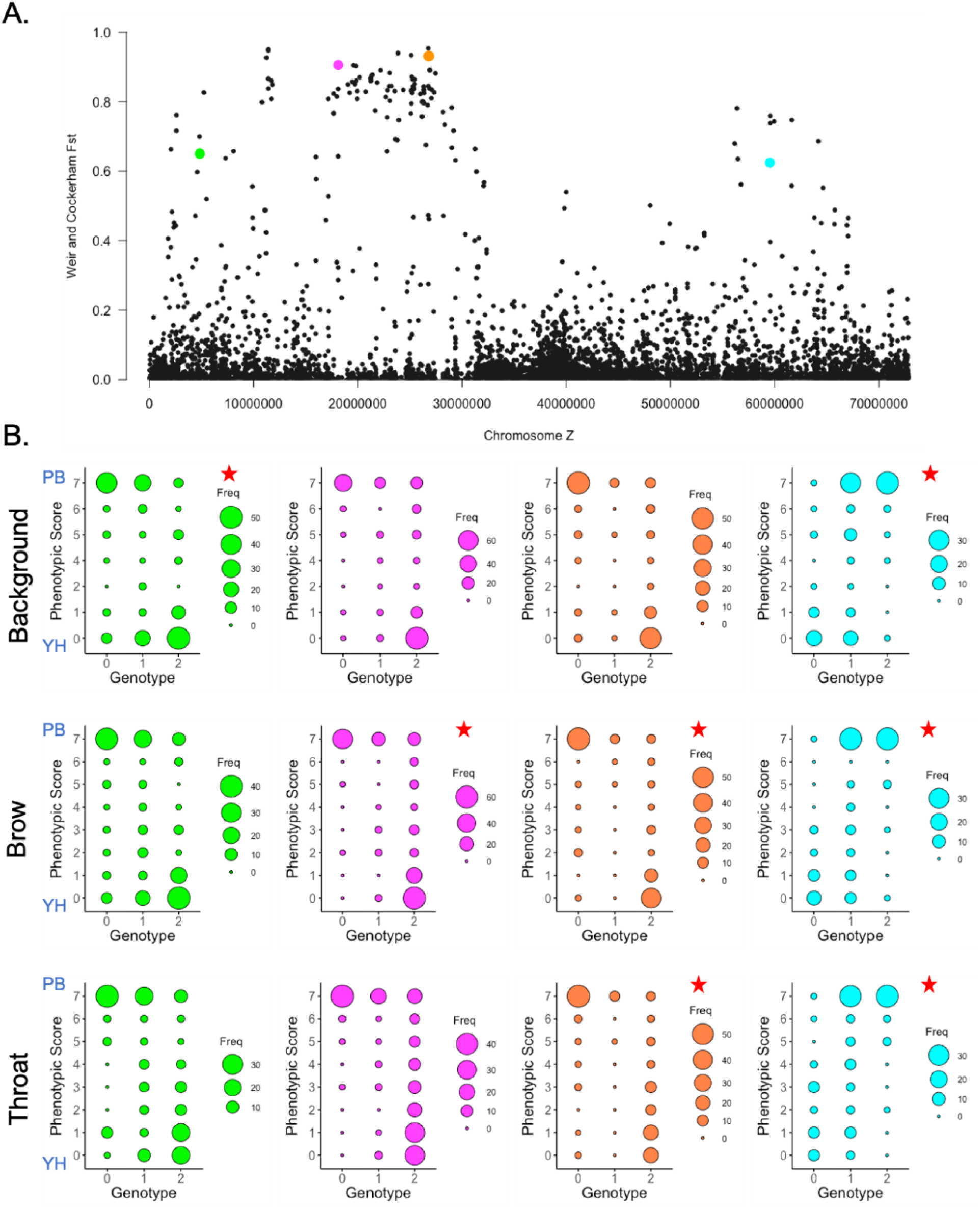
Investigation of potential dominance interactions between the alleles of SNPs that are significantly associated with plumage variation as identified using GEMMA. **A)** Relative differentiation (*F*ST) of 11,147 SNPs on the Z chromosome calculated based on comparisons between allopatric yellowhammers (n = 53) and allopatric pine buntings (n = 42). Four SNPs are highlighted in this panel: Z.4835388 (green), Z.18131016 (magenta), Z.26812248 (orange) and Z.59571043 (blue). **B)** Balloon plots illustrating the number of individuals possessing a particular phenotypic score at one of three plumage traits with a particular genotype at one of the four SNPs highlighted in panel A. The plumage traits considered are: the background plumage colour (“Background”), the amount of chestnut at the brow (“Brow”) and the amount of chestnut at the throat (“Throat”). The SNP identities are indicated with the same colours as in panel A: Z.4835388 (green), Z.18131016 (magenta), Z.26812248 (orange) and Z.59571043 (blue). Dot size in each balloon plot indicates the number of individuals possessing a specific phenotypic score-genotype combination. Red stars indicate whether each of the four highlighted SNPs is significantly associated with phenotypic variation at the plumage trait of interest as determined using GEMMA. Blue labels “YH” (Yellowhammer) and “PB” (Pine Bunting) indicate phenotypic scores more commonly associated with “pure” members of each taxon.

The SNP Z.4835388—located to the left of the putative inversion (Figure 7A)—was significantly associated with background colour and the genomic region surrounding it was weakly associated with facial plumage traits (Table 2; Figure 6). Comparisons of phenotypic scores against Z.4835388 genotypes did not suggest dominance at any of the traits as heterozygotes did not show a strong bias towards one ancestral plumage phenotype and both homozygotes possessed variable plumage phenotypes (Figure 7B).

The SNP Z.18131016—located near the beginning of the putative inversion (Figure 7A)— was significantly associated with brow plumage variation (Table 3; Figure 6B). The genomic region surrounding this SNP was also moderately associated with background colour and throat plumage (Figure 6A, Supplementary Figure 6C). For Z.18131016, we saw patterns consistent with dominance as the heterozygous and one homozygous genotype showed a strong bias towards scores of “7”—the pure pine bunting phenotype—for all plumage traits (Figure 7B). The remaining homozygous genotype was associated with variable phenotypic scores for all three traits including those associated with phenotypic yellowhammers and hybrids. These results suggest that the putative “pine bunting” allele at this SNP is dominant over the putative “yellowhammer” allele in its effects on plumage traits.

The SNP Z.26812248—located near the end of the putative inversion (Figure 7A)— was significantly associated with brow and throat plumage (Table 3-4; Figure 6B-C). The genomic region surrounding this SNP was also moderately associated with variation in background colour (Figure 6A). As seen for SNP Z.18131016, comparisons of phenotypic scores and Z.26812248 genotypes showed patterns consistent with a putative “pine bunting” allele being dominant over a putative “yellowhammer” allele for all three plumage traits (Figure 7B). The heterozygous and one homozygous genotype were highly associated with pure pine bunting phenotypic scores while the other homozygous genotype was associated with variable phenotypic scores.

Finally, the SNP Z.59571043—located to the right of the putative inversion (Figure 7A)—was significantly associated with all three plumage traits (Table 2-4; Figure 6). Despite these associations, heterozygous and homozygous genotypes did not show any bias towards a particular ancestral phenotype for any of the plumage traits suggesting an absence of dominance interactions at this SNP (Figure 7B).

## Discussion

The yellowhammer and pine bunting have historically been considered separate species because of their highly divergent plumage phenotypes (Panov et al. 2003; Rubtsov and Tarasov, 2017), and clear nuclear genetic differentiation between their allopatric populations (Irwin et al. 2009; Nikelski et al. 2023) might be viewed as supporting this treatment. However, widespread hybridization in sympatry indicated by several intermediate plumage phenotypes (Panov et al. 2003; 2007; Rubtsov, 2007; Rubtsov & Tarasov, 2017) as well as evidence of mitochondrial DNA (Irwin et al. 2009) and mitonuclear gene introgression (Nikelski et al. 2023) suggest that reproductive isolation is weak and that yellowhammers and pine buntings might best be considered different forms of a single species. Altogether, these conflicting observations suggest two possibilities for the future of this system: further population divergence or population merging. Our work exploring patterns of genetic differentiation and hybridization between yellowhammers and pine buntings across Eurasia suggests that these taxa are on the cusp of these two trajectories. Here, we confirmed the presence of strong genetic differentiation between allopatric yellowhammers and pine buntings (Irwin et al. 2009; Nikelski et al. 2023) that extends into what we classify as the near-sympatric zone, but we also found higher than expected levels of genetic admixture in the sympatric region suggesting that reproductive barriers between taxa are weak.

The extensive hybridization documented between yellowhammers and pine buntings over decades of research (Panov et al. 2003; 2007; Rubtsov, 2007; Rubtsov & Tarasov, 2017) was exemplified in our genomic results showing high genetic admixture within phenotypic hybrids and, surprisingly, moderate genetic admixture within phenotypically pure sympatric individuals with this latter pattern being more prominent in sympatric pine buntings. These trends were maintained when accounting for the phenotypic class of sympatric individuals, as genetic ancestry from both yellowhammers and pine buntings was represented in all plumage types. This prevalence of sympatric individuals with admixed ancestry implies an absence of strong pre-zygotic barriers that prevent interbreeding between yellowhammers and pine buntings. Supporting this idea, past surveys of sympatric yellowhammer and pine bunting male mating territories found that individuals were equally territorial to con-and hetero-specifics (Rubtsov & Tarasov, 2017). If we assume that male territorial response can be used as a proxy for female mate choice as has been done in other avian systems (e.g. Balakrishnan & Sorenson, 2006; Uy et al. 2009; Cruz-Yepez et al. 2020), this finding suggests that sympatric females may be non-discriminatory or only weakly discriminatory when choosing a mate. On the other hand, past surveys of the yellowhammer and pine bunting hybrid zone reported a potential tendency towards assortative mating among individuals (Rubtsov, 2022). The relationship between phenotypic class and genomic variation within the sympatric zone suggests that assortative mating via plumage traits could, in fact, act as a pre-zygotic barrier between yellowhammers and pine buntings as “PC” and “PL” individuals possess distinct genotypes on average. However, if such a barrier were to exist, it is likely that it would become increasingly ineffective as the presence of sympatric individuals with mismatched genotypes and plumage phenotypes (i.e. phenotypically admixed individuals with putatively pure genotypes and phenotypically pure individuals with putatively admixed genotypes) suggests that these two factors are becoming disassociated with time.

Our genomic data showed that there are not distinct yellowhammer and pine bunting gene pools within the sympatric zone. This result is consistent with relatively uninhibited gene flow between sympatric populations and a lack of strong post-zygotic barriers within the system. Indeed, the degree and variation of genetic admixture seen in the sympatric zone implies that hybrids are not only viable, as was confirmed in previous research (Lohrl, 1967 cited in Panov et al., 2003), but that they are also fertile and likely possess few deficiencies that impact their fitness. However, life duration data from the yellowhammer and pine bunting sympatric zone provides tentative support for the life expectancies of individuals in the “SC”, “CH”, “YH”, “WH”, “LH” and “SL” phenotypic classes being slightly reduced compared to those of individuals in the “PC” and “PL” phenotypic classes (Rubtsov, 2022). This survival bias implies that there is some amount of post-zygotic isolation between sympatric “PC” and “PL” individuals—which are representative of yellowhammers and pine buntings respectively in their purest genomic forms—which could prevent complete merging of these two taxa. Nevertheless, our genomic data suggests that any post-zygotic barriers between yellowhammers and pine buntings (or between “PC” and “PL” individuals) are not strong enough to prevent extensive genetic admixture within the sympatric zone. This assertion coupled with the identification of sympatric individuals with mismatched plumage phenotypes and genomic ancestries (including “PC” and “PL” individuals) makes connecting Rubtsov’s (2022) findings to our own very complex. In any case, the state of post-zygotic isolation in the yellowhammer and pine bunting system warrants further investigation.

In the face of extensive hybridization, phenotypic diversity may be reduced due to genetic homogenization and the loss of distinct haplotypes that are responsible for phenotypic differences between populations. However, diversity can be maintained within a system when there are genomic structures that suppress recombination and maintain linkage disequilibrium between the genetic loci that underlie phenotypes (Nachman, 2002). Chromosomal inversions are an example of such structures as different orientations of an inversion usually have greatly reduced recombination allowing them to maintain linkage disequilibrium and accumulate genetic differentiation (Kirkpatrick, 2010). In some cases, these highly differentiated, linked haploblocks can underlie important reproductive barriers between taxa (Noor et al. 2001; Rieseberg, 2001), but, in others, the two forms of the inversion can segregate within the system maintaining elevated levels of genetic—and consequently phenotypic—diversity among interbreeding individuals (Tuttle et al. 2016; Brelsford et al. 2020).

In the yellowhammer and pine bunting system, we found that genetic variation at a previously identified Z chromosome “island of differentiation” (Nikelski et al. 2023) separated male individuals into three distinct genetic clusters. More specifically, the intermediate male cluster could be attributed to a large block of heterozygosity within this differentiation island and this finding was supported by an analysis of linkage disequilibrium across chromosome Z. Altogether, these results suggest that the identified differentiation island is an area of low recombination that has produced two highly linked, genetically divergent haploblocks. It is highly likely that these divergent blocks represent different orientations of a chromosomal inversion that are segregating within the system, suggesting that intermediate males inherited one copy of each form of the inversion and members of the two remaining male clusters inherited two copies of the same inversion form. This putative chromosomal inversion acts as a polymorphism of the system and contributes genetic and—based on its association with plumage traits discussed below—phenotypic diversity among individuals.

Yellowhammers and pine buntings display highly divergent phenotypes in a number of plumage traits. These characteristics were crucial when granting them species designations and have been considered the most likely candidates for reproductive barriers in the system (Panov et al. 2003; Rubtsov & Tarasov, 2017). Yet, our genomic results show some discordance between individual phenotype and genotype within the sympatric zone implying that plumage is not necessarily a clear indicator of genetic ancestry. When we conducted admixture mapping to associate plumage variation with genetic variation across the system, we found that all three divergent plumage traits—background colour, amount of chestnut at the brow and amount of chestnut at the throat—were strongly correlated with loci on the Z chromosome with the latter two characteristics being specifically correlated with the putative inversion polymorphism.

In addition to reporting a strong association between the putative chromosome Z inversion polymorphism and plumage, we also showed evidence that the “pine bunting” inversion form acts dominantly to the “yellowhammer” inversion form in its effects on the plumage traits studied here. This dominance pattern was observed for two SNPs positioned within either end of the inversion in terms of their association with all three plumage traits. In contrast, SNPs located outside of the inversion but that were significantly associated with plumage variation did not show evidence of dominance interactions. SNPs on other chromosomes may also affect phenotypic variation in this system, and are likely responsible for some of the unexplained variation seen in our dominance analysis.

If the “pine bunting” form of the inversion is dominant over the “yellowhammer” form in its effects on plumage traits, this relationship may explain why sympatric yellowhammers show less overall genomic admixture compared to sympatric pine buntings. In such a dominance framework, individuals who are homozygous or heterozygous for the “pine bunting” form of the inversion would display a pine bunting phenotype despite potentially high ancestry from yellowhammers at other genomic regions. As such, admixture is masked by dominance and individuals appear as phenotypic pine buntings. For individuals who are homozygous for the “yellowhammer” form of the inversion, admixture would not be masked and could potentially influence the plumage phenotype. Here, individuals who possess relatively pure yellowhammer ancestry in addition to two versions of the “yellowhammer” inversion form would display a pure yellowhammer phenotype while individuals who possess two versions of the “yellowhammer” inversion and some admixture from pine buntings would display a hybrid phenotype.

In terms of the specific genes responsible for plumage variation in this system, we found that the SNP most significantly associated with each of the three plumage traits did not occur within or near an annotated gene known to be related to colouration or plumage. However, additional SNPs significantly associated with plumage variation were located within genes that could mediate plumage phenotypes.

For background plumage colour, we found a highly significant association with the gene ABCA1 (ATP Binding Cassette Subfamily A Member 1) on chromosome Z. The ABCA1 gene is involved in the translocation of phospholipids across membranes as well as the formation of high-density lipoproteins (HDLs; reviewed in Oram & Vaughan, 2000) that are used to transport carotenoids around the body (Clevidence & Bieri, 1993). A mutation in ABCA1 among Wisconsin hypoalpha mutant (WHAM) chickens is responsible for low levels of HDLs and, as a result, low levels of carotenoids in chicken tissues (Connor et al. 2007), producing a phenotype of a white beak and skin versus a yellow beak and skin as well as colourless plasma (Attie et al. 2002). These results show how ABCA1 gene mutations can have a direct effect on carotenoid transport and cause a shift in colour similar to what is seen in our system, suggesting that it is a strong candidate for regulation of plumage patterning between yellowhammers and pine buntings. In addition to ABCA1, variation in background plumage colour was also significantly associated with CHD6 (Chromodomain Helicase Binding Protein 6) on chromosome 20. Currently, we are not aware of published connections between CHD6 and the regulation of either plumage patterning or colouration.

Moving onto facial plumage patterns, we noted a tight correspondence between the loci significantly associated with brow and throat plumage as was hypothesized previously in the yellowhammer and pine bunting system (Panov et al. 2003). Only one gene was significantly associated with both plumage traits: GRAMD3 (GRAM domain containing 3). Little is known about the function of GRAMD3, but it is highly associated with retinal pigment epithelial cells in humans (Strunnikova et al. 2010). More notably, GRAMD3 has been proposed as a candidate gene for the *Id* locus (Xu et al. 2017). The *Id* locus controls dermal shank pigmentation in chickens which is a trait defined by the accumulation of melanin in the dermis of the leg (McGibbon, 1974). This connection between GRAMD3 and melanin colouration is consistent with the presumed melanin-based brow and throat patterning in the yellowhammer and pine bunting system, indicating that this gene may be important in the regulation of these traits. Other potentially important genes associated with variation in brow plumage included: FER (FER tyrosine kinase), which is associated with melanoma in humans (Ivanova et al. 2019); EFNA5 (Ephrin A5), which is associated with piebald skin pigmentation in sheep (García-Gámez et al. 2011); and APC (APC regulator of Wnt signaling pathway), which is associated with feather development (Widelitz et al. 2000). The CEP120 (Centrosomal protein 120), PJA2 (Praja ring finger ubiquitin ligase 2), MCC (MCC regulator of Wnt signaling pathway) and ECPAS (ECM29 proteasome adapter and scaffold) genes were also associated with brow plumage, but their known functions have not been linked to colouration, plumage patterning or feather development. Although it is possible that these additional genes play a role in brow plumage patterning, it should be noted that all of them were found within the putative inversion meaning that the observed associations may be due to linkage with a specific gene, like GRAMD3, within the inversion that is regulating colouration patterns.

To summarize, allopatric and near-sympatric populations of yellowhammers and pine buntings form genetically discrete clusters well separated from each other. However, due to an apparent lack of strong reproductive barriers, this relationship breaks down in sympatry where hybridization and genetic admixture is extensive. Much of the genetic differentiation seen between allopatric and near-sympatric yellowhammers and pine buntings is driven by a large “island of differentiation” on the Z chromosome which appears to house an inversion polymorphism. This putative inversion is associated with the highly divergent plumage traits currently used to distinguish between the taxa (Panov et al. 2003; Rubtsov & Tarasov, 2017) and is also subject to dominance interactions with the “pine bunting” form of the inversion being dominant over the “yellowhammer” form in its effects on plumage traits. The different inversion forms generally follow species designations in allopatry and near-sympatry, but may be beginning to flow between taxa in sympatry much like the rest of the nuclear genome. Nevertheless, the non-recombining nature of the putative inversion polymorphism hints that it could continue to contribute towards plumage variation within this system in the future.

Keeping this final point in mind, these findings provide an interesting narrative for the evolution of the yellowhammer and pine bunting system. During the Pleistocene glaciations, separated populations appear to have diverged genetically while in allopatric isolation. This divergence included the evolution of an apparent inversion polymorphism that included loci controlling plumage traits. Eventually, the accumulation of mutations within the putative inversion produced highly divergent plumage phenotypes between what are now be classified as yellowhammers and pine buntings. Yet, genetic differentiation within and outside the inversion was not great enough to constitute strong reproductive barriers between taxa, resulting in widespread hybridization and gene flow upon secondary contact. While the mitochondrial genome (Irwin et al. 2009) and a large part of the nuclear genome have become genetically homogenized between yellowhammers and pine buntings, the putative inversion on chromosome Z has resisted homogenization allowing for the retention of ancestral plumage phenotypes alongside hybrid phenotypes within the sympatric zone despite the majority of individuals being genetically admixed. As well, possible weak post-zygotic isolation between sympatric individuals possessing “PC” and “PL” phenotypes (Rubtsov, 2022) may allow for the continued retention of some genetic differentiation between yellowhammers and pine buntings in the sympatric zone in spite of hybridization and gene flow.

Looking forward, the yellowhammer and pine bunting system faces two possible futures. If proposed post-zygotic isolation between sympatric “PC” and “PL” individuals (Rubtsov, 2022) is strong enough or evolves to be strong enough to maintain distinct yellowhammer and pine bunting genotypes, time may see a decline in hybrid populations within the sympatric zone due to lower hybrid survival (Coyne & Orr, 2004; Price, 2008). Such fitness consequences could drive the evolution of pre-zygotic barriers, likely assortative mating based on plumage traits, through the process of reinforcement (Butlin, 1987). Eventually, interbreeding would cease entirely between sympatric yellowhammers and pine buntings and speciation between these taxa would be complete (Coyne & Orr, 2004; Price, 2008). Alternatively, if post-zygotic barriers between populations remain weak, the area of sympatry and hybridization between yellowhammers and pine buntings may continue to expand (Panov et al. 2003; Rubtsov & Tarasov, 2017) potentially until the two taxa merge into one broadly sympatric and freely interbreeding population. Yet, even in the face of population merging, it is likely that some version of the current yellowhammer and pine bunting plumage patterns will be preserved as a conspicuous within-species polymorphism. This is due to the protection afforded to the plumage-controlling haplotypes by their apparent position within a chromosomal inversion and to potential weak post-zygotic barriers between sympatric “PC” and “PL” individuals (Rubtsov, 2022).

Extreme phenotypic variation within a system of interbreeding populations has only been observed for a select number of avian taxa including the white-throated sparrows (*Zonotrichia albicollis*; Tuttle, 2003), the Gouldian finches (*Chloebia gouldiae*; Southern 1945; Brush & Seifried, 1968) and the ruffs (*Calidris pugnax*; Hogan-Warburg, 1966; Jukema & Piersma, 2006). Thus, in its relative novelty, the high phenotypic diversity seen among yellowhammers and pine buntings emphasizes the role of hybridization as both a destructive and creative force in evolution. Though hybridization is homogenizing the genomes of two divergent taxa, it is also creating a system or—in the case of population merging—a single population of interbreeding organisms with a large amount of phenotypic variation. In doing this, hybridization has the potential to inflate a system/population’s evolvability to a wider range of environmental conditions (Parsons et al. 2011; Grant & Grant 2019; Kagawa & Seehausen, 2020). This higher evolvability can safeguard the system/population from extinction and may even pave the way for continued divergence and speciation if evolutionary pressures change. Such implications of destructive and constructive hybridization warrant further discussion—in regard to the yellowhammers and pine bunting system and others—as researchers continue to investigate the evolutionary forces mediating genetic differentiation and develop a more holistic understanding of the speciation process.

### Author Contributions

E. N., D.I., and A.S.R. designed this study. A.S.R. obtained blood and tissue samples. E. N. and A.S.R. performed molecular procedures. E. N. completed bioinformatic and statistical analyses and wrote this manuscript with contributions from D.I. and A.S.R.

## Acknowledgements

We would like to acknowledge Dolph Schluter, Eric Taylor, Judith Mank, Elizabeth Natola, Rashika Ranasinghe, Kenneth Askelson, Finola Fogarty, Quinn McCallum, Ana Barreira, Jamie Clarke, Armando Geraldes and Jessica Irwin for their invaluable comments and advice over the course of this study. We would also like to thank Madelyn Ore and the Tazeev family for their assistance during fieldwork. Finally, we wish to recognize The Bell Museum, The Burke Museum of Natural History and Culture, The Field Museum, The State Darwin Museum, The Swedish Museum of Natural History, The Zoological Museum of the Zoological Institute of the Russian Academy of Sciences, and the Zoological Museum of the University of Copenhagen for providing us with additional samples that were utilized in this work. Research funding was provided by the Natural Sciences and Engineering Research Council of Canada (NSERC CGSM award to E.N., Discovery Grants RGPIN-2017-03919 and RGPAS-2017-507830 awarded to D.I.) and by the Werner and Hildegard Hesse research awards (Research award in Ornithology and Fellowship in Ornithology awarded to E.N. by the University of British Columbia).

## Data Accessibility Statement

The raw DNA reads utilized in this study are available on the NBCI Sequence Read Archive (BioProject PRJNA768601). Read processing codes, barcodes and genotype data are available on a Dryad repository associated with a previous publication (Nikelski et al. 2023; https://doi.org/10.5061/dryad.tmpg4f538). R codes associated with the genetic and statistical analyses performed in this study will be made available in a separate Dryad repository upon publication.

## References

1. Andrade, P., Pinho, C., de Lanuza, G. P., Afonso, S., Brejcha, J., Rubin, C., Wallerman, O., Pereira, P., Sabatino, S. J., Bellati, A., Pellitteri-Rosa, D., Bosakova, Z., Bunikis, I., Carretero, M. A., Feiner, N., Marsik, P., Paupério, F., Salvi, D., Soler, L., … Carneiro, M. (2019). Regulatory changes in pterin and carotenoid genes underlie balanced color polymorphisms in the wall lizard. Proceedings of the National Academy of Sciences, 116(12), 5633–5642. https://doi:10.1073/pnas.1820320116

2. Alexander, D. H., Novembre, J., & Lange, K. (2009). Fast model-based estimation of ancestry in unrelated individuals. Genome Research, 19(9), 1655–1664. https://doi:10.1101/gr.094052.109

3. Attie, A. D., Hamon, Y., Brooks-Wilson, A. R., Gray-Keller, M. P., MacDonald, M. L. E., Rigot, V., Tebon, A., Zhang, L-H., Mulligan, J. D., Singaraja, R. R., Bitgood, J. J., Cook, M. E., Kastelein, J. J. P., Chimini, G., & Hayden, M. R. (2002). Identification and functional analysis of a naturally occurring E89K mutation in the ABCA1 gene of the WHAM chicken. Journal of Lipid Research, 43(10), 1610–1617. https://doi:10.1194/jlr.M200223-JLR200

4. Ayala, D., Guerrero, R. F., & Kirkpatrick, M. (2013). Reproductive isolation and local adaptation quantified for a chromosome inversion in a malaria mosquito. Evolution, 67(4), 946–958. https://doi:10.1111/j.1558-5646.2012.01836.x

5. Balakrishnan, C. N., & Sorenson, M. D. (2006). Song discrimination suggests premating isolation among sympatric indigobird species and host races. Behavioral Ecology, 17(3), 473–478. https://doi:10.1093/beheco/arj052

6. Barton, N. H., & Hewitt, G. M. (1985). Analysis of hybrid zones. Annual Review of Ecology and Systematics, 16(1), 113–148. https://doi:10.1146/annurev.es.16.110185.000553

7. Barton, N. H., & Hewitt, G. M. (1989). Adaptation, speciation and hybrid zones. Nature, 341(6242), 497–503. https://doi:10.1038/341497a0

8. Bateson, W. 1909. Heredity and variation in modern lights. In A. C. Seward (Eds.) Darwin and Modern Science (pp. 85–101). Cambridge University Press.

9. Blair, W. F. (1955). Mating call and stage of speciation in the Microhyla olivacea-M. carolinensis complex. Evolution, 9(4), 469–480. https://doi:10.1111/j.1558-5646.1955.tb01556.x

10. Brelsford, A., Toews, D. P. L., & Irwin, D. E. (2017). Admixture mapping in a hybrid zone reveals loci associated with avian feather coloration. Proceedings of the Royal Society B: Biological Sciences, 284(1866), 1–9. https://doi:10.1098/rspb.2017.1106

11. Brelsford, A., Purcell, J., Avril, A., Tran Van, P., Zhang, J., Brütsch, T., Sundström, L., Helanterä, H., &Chapuisat, M. (2020). An ancient and eroded social supergene is widespread across Formica ants. Current Biology, 30(2), 304–311. https://doi:10.1016/j.cub.2019.11.032

12. Bridle, J. R., Saldamando, C. I., Koning, W., & Butlin, R. K. (2006). Assortative preferences and discrimination by females against hybrid male song in the grasshoppers Chorthippus brunneus and Chorthippus jacobsi (Orthoptera: Acrididae). Journal of Evolutionary Biology, 19(4), 1248–1256. https://doi:10.1111/j.1420-9101.2006.01080.x

13. Brush, A. H., & Seifried, H. (1968). Pigmentation and feather structure in genetic variants of the Gouldian finch, Poephila gouldiae. The Auk, 85(3), 416–430. https://doi:10.2307/4083290

14. Butlin, R. (1987). Speciation by reinforcement. Trends in Ecology & Evolution, 2(1), 8–13. https://doi:10.1016/0169-5347(87)90193-5

15. Cerdá-Reverter, J. M., Haitina, T., Schiöth, H. B., & Peter, R. E. (2005). Gene structure of the goldfish agouti-signaling protein: A putative role in the dorsal-ventral pigment pattern of fish. Endocrinology, 146(3), 1597–1610. https://doi:10.1210/en.2004-1346

16. Chang, C. C., Chow, C. C., Tellier, L. C., Vattikuti, S., Purcell, S. M., & Lee, J. J. (2015). Second-generation PLINK: Rising to the challenge of larger and richer datasets. Gigascience, 4(1). https://doi:10.1186/s13742-015-0047-8

17. Clevidence, B. A., & Bieri, J. G. (1993). Association of carotenoids with human plasma lipoproteins. Methods in Enzymology, 214, 33–46.

18. Connor, W. E., Duell, P. B., Kean, R., & Wang, Y. (2007). The prime role of HDL to transport lutein into the retina: Evidence from HDL-deficient WHAM chicks having a mutant ABCA1 transporter. Investigative Ophthalmology & Visual Science, 48(9), 4226–4231. https://doi:10.1167/iovs.06-1275

19. Coyne, J. A., & Orr, H. A. (2004). Speciation. Sinauer Associates.

20. Cruz-Yepez, N., González, C., & Ornelas, J. F. (2020). Vocal recognition suggests premating isolation between lineages of a lekking hummingbird. Behavioral Ecology, 31(4), 1046–1053. https://doi:10.1093/beheco/araa050

21. Danecek, P., Auton, A., Abecasis, G., Albers, C. A., Banks, E., DePristo, M. A., Handsaker, R. E., Lunter, G., Marth, G. T., Sherry, S. T., McVean, G., Durbin, R. & 1000 Genomes Project Analysis Group. (2011). The variant call format and VCFtools. Bioinformatics, 27(15), 2156–2158. https://doi:10.1093/bioinformatics/btr330

22. Danley, P. D., DeCarvalho, T. N., Fergus, D. J., & Shaw, K. L. (2007). Reproductive asynchrony and the divergence of Hawaiian crickets. Ethology, 113(12), 1125–1132. https://doi:10.1111/j.1439-0310.2007.01430.x

23. Dobzhansky, T. 1937. Genetics and the Origin of Species. Columbia University Press. Dobzhansky, T. (1940). Speciation as a stage in evolutionary divergence. The American Naturalist, 74(753), 312–321. https://doi:10.1086/280899

24. Edwards, S. V., Kingan, S. B., Calkins, J. D., Balakrishnan, C. N., Jennings, W. B., Swanson, W. J., & Sorenson, M. D. (2005). Speciation in birds: Genes, geography, and sexual selection. Proceedings of the National Academy of Sciences, 102(Suppl 1), 6550–6557. https://doi:10.1073/pnas.0501846102

25. Elshire, R. J., Glaubitz, J. C., Sun, Q., Poland, J. A., Kawamoto, K., Buckler, E. S., & Mitchell, S. E. (2011). A robust, simple genotyping-by-sequencing (GBS) approach for high diversity species. PloS One, 6(5), e19379. https://doi:10.1371/journal.pone.0019379

26. Fitzpatrick, B. M. (2012). Estimating ancestry and heterozygosity of hybrids using molecular markers. BMC Evolutionary Biology, 12(1), 1–14. https://doi:10.1186/1471-2148-12-131

27. García-Gámez, E., Reverter, A., Whan, V., McWilliam, S. M., Arranz, J. J., Kijas, J., & International Sheep Genomics Consortium. (2011). Using regulatory and epistatic networks to extend the findings of a genome scan: Identifying the gene drivers of pigmentation in merino sheep. PloS One, 6(6), e21158–e21158. https://doi:10.1371/journal.pone.0021158

28. Gompert, Z., Mandeville, E. G., & Buerkle, C. A. (2017). Analysis of population genomic data from hybrid zones. Annual Review of Ecology, Evolution, and Systematics, 48(1), 207–229. https://doi:10.1146/annurev-ecolsys-110316-022652

29. Grant, P. R., & Grant, B. R. (1997). Genetics and the origin of bird species. Proceedings of the National Academy of Sciences, 94(15), 7768–7775. https://doi:10.1073/pnas.94.15.7768

30. Grant, P. R., & Grant, B. R. (2019). Hybridization increases population variation during adaptive radiation. PNAS, 116(46), 23216–23224. https://doi:10.1073/pnas.1913534116

31. Gross, J. B., Borowsky, R., & Tabin, C. J. (2009). A novel role for Mc1r in the parallel evolution of depigmentation in independent populations of the cavefish Astyanax mexicanus. PLoS Genetics, 5(1), e1000326–e1000326. https://doi:10.1371/journal.pgen.1000326

32. Kagawa, K., & Seehausen, O. (2020). The propagation of admixture-derived adaptive radiation potential. Proceedings of the Royal Society B: Biological Sciences, 287(1934), 20200941–20200941. https://doi:10.1098/rspb.2020.0941

33. Haupaix, N., Curantz, C., Bailleul, R., Beck, S., Robic, A., & Manceau, M. (2018). The periodic coloration in birds forms through a prepattern of somite origin. Science, 361(6408), 1216. https://doi:10.1126/science.aar4777

34. Hedrick, P. W., Smith, D. W., & Stahler, D. R. (2016). Negative-assortative mating for color in wolves. Evolution, 70(4), 757–766. https://doi:10.1111/evo.12906

35. Hewitt, G. M. (1988). Hybrid zones-natural laboratories for evolutionary studies. Trends in Ecology & Evolution, 3(7), 158–167. https://doi:10.1016/0169-5347(88)90033-x

36. Hogan-Warburg, A. J. (1966). Social behavior of the ruff, Philomachus pugnax (L.) (Vol. 54). Brill Archive.

37. Hubbard, J. K., Uy, J. A. C., Hauber, M. E., Hoekstra, H. E., & Safran, R. J. (2010). Vertebrate pigmentation: From underlying genes to adaptive function. Trends in Genetics, 26(5), 231–239. https://doi:10.1016/j.tig.2010.02.002

38. Irwin, D. E. (2020). Assortative mating in hybrid zones is remarkably ineffective in promoting speciation. The American Naturalist, 195(6), E150–E167. https://doi.org/10.1086/708529

39. Irwin, D. E., Milá, B., Toews, D. P. L., Brelsford, A., Kenyon, H. L., Porter, A. N., Grossen, C., Delmore, K. E., Alcaide, M., & Irwin, J. H. (2018). A comparison of genomic islands of differentiation across three young avian species pairs. Molecular Ecology, 27(23), 4839–4855. https://doi:10.1111/mec.14858

40. Irwin, D. E., Rubtsov, A. S., & Panov, E. N. (2009). Mitochondrial introgression and replacement between yellowhammers (Emberiza citrinella) and pine buntings (Emberiza leucocephalos) (Aves: Passeriformes). Biological Journal of the Linnean Society, 98(2), 422–438. https://doi:10.1111/j.1095-8312.2009.01282.x

41. Ivanova, I., Arulanantham, S., Barr, K., Cepeda, M., Parkins, K., Hamilton, A., Johnson, D., Penuela, S., Hess, D. A., Ronald, J. A., & Dagnino, L. (2019). Targeting FER kinase inhibits melanoma growth and metastasis. Cancers, 11(3), 419. https://doi:10.3390/cancers11030419

42. Jukema, J., & Piersma, T. (2006). Permanent female mimics in a lekking shorebird. Biology Letters, 2(2), 161–164. https://doi:10.1098/rsbl.2005.0416

43. Kassambara, A. (2017). ggpubr: Publication Ready Plots. Statistical Tools for High-Throughput Data Analysis. http://www.sthda.com/english/articles/24-ggpubr-publication-ready-plots/

44. Kingsley, E. P., Manceau, M., Wiley, C. D., & Hoekstra, H. E. (2009). Melanism in Peromyscus is caused by independent mutations in agouti. PloS One, 4(7), e6435–e6435. https://doi:10.1371/journal.pone.0006435

45. Kirkpatrick, M. (2010). How and why chromosome inversions evolve. PLoS Biology, 8(9), e1000501. https://doi:10.1371/journal.pbio.1000501

46. Lohrl, H. (1967). Zur Verwandtschaft von Fichtenammer (Emberiza leucocephalos) und Goldammer (Emberiza citrinella). Vogelwelt, 88, 148–152.

47. Lowry, D. B., & Willis, J. H. (2010). A widespread chromosomal inversion polymorphism contributes to a major life-history transition, local adaptation, and reproductive isolation. PLoS Biology, 8(9), e1000500. https://doi:10.1371/journal.pbio.1000500

48. Manichaikul, A., Mychaleckyj, J. C., Rich, S. S., Daly, K., Sale, M., & Chen, W. (2010). Robust relationship inference in genome-wide association studies. Bioinformatics, 26(22), 2867–2873. https://doi:10.1093/bioinformatics/btq559

49. Mason, N. A., & Bowie, R. C. K. (2020). Plumage patterns: Ecological functions, evolutionary origins, and advances in quantification. The Auk, 137(4), 1–29. https://doi.org/10.1093/auk/ukaa060

50. Mayr, E. 1942. Systematics and the Origin of Species. Columbia University Press, New York. McGibbon, W. H. (1974). A shank color mutation in Cornell random bred S.C. white leghorns. Poultry Science, 53(3), 1251–1253. https://doi:10.3382/ps.0531251

51. Moore, I. T., Bonier, F., & Wingfield, J. C. (2005). Reproductive asynchrony and population divergence between two tropical bird populations. Behavioral Ecology, 16(4), 755–762. https://doi:10.1093/beheco/ari049

52. Muller, H. J. (1942). Isolating mechanisms, evolution, and temperature. In Biology Symposium (Vol. 6, pp. 71-125).

53. Nachman, M. (2002). Variation in recombination rate across the genome: evidence and implications. Current Opinion in Genetics & Development, 12(6), 657–663. https://doi:10.1016/s0959-437x(02)00358-1

54. Nikelski, E., Rubtsov, A. S., & Irwin, D. (2023). High heterogeneity in genomic differentiation between phenotypically divergent songbirds: A test of mitonuclear co-introgression. Heredity, 130(1), 1–13. https://doi:10.1038/s41437-022-00580-8

55. Noor, M. A. F., Grams, K. L., Bertucci, L. A., & Reiland, J. (2001). Chromosomal inversions and the reproductive isolation of species. Proceedings of the National Academy of Sciences, 98(21), 12084–12088. https://doi:10.1073/pnas.221274498

56. Oram, J. F., & Vaughan, A. M. (2000). ABCA1-mediated transport of cellular cholesterol and phospholipids to HDL apolipoproteins. Current Opinion in Lipidology, 11(3), 253–260. https://doi:10.1097/00041433-200006000-00005

57. Ortiz-Barrientos, D., Engelstädter, J., & Rieseberg, L. H. (2016). Recombination rate evolution and the origin of species. Trends in Ecology & Evolution, 31(3), 226–236. https://doi:10.1016/j.tree.2015.12.016

58. Panov, E. N., Rubtsov, A. S., & Monzikov, D. G. (2003). Hybridization between yellowhammer and pine bunting in Russia. Dutch Birding, 25, 17–31.

59. Panov, E. N., Rubtsov, A. S., & Mordkovich, M. V. (2007). New data on the relationships between two species of buntings (Emberiza citrinella and E. leucocephalos) hybridizing in the areas of overlap of their ranges, Zoologicheskii Zhurnal, 86(11), 1362–1378.

60. Parsons, K. J., Son, Y. H., & Craig Albertson, R. (2011). Hybridization promotes evolvability in African cichlids: Connections between transgressive segregation and phenotypic integration. Evolutionary Biology, 38(3), 306–315. https://doi:10.1007/s11692-011-9126-

61. Price, T. (2008). Speciation in birds. Roberts and Co.R Core Team (2014). R: A language and environment for statistical computing. Vienna, Austria: R Foundation for Statistical Computing. Retrieved from http://www.r-project.org/

62. Rieseberg, L. H. (2001). Chromosomal rearrangements and speciation. Trends in Ecology & Evolution, 16(7), 351–358. https://doi:10.1016/s0169-5347(01)02187-5

63. Rosenblum, E. B., Hoekstra, H. E., & Nachman, M. W. (2004). Adaptive reptile color variation and the evolution of the Mc1r gene. Evolution, 58(8), 1794–1808. https://doi.org/10.1111/j.0014-3820.2004.tb00462.x

64. Rubtsov, A.S. (2007). Variability of songs of the yellowhammer (Emberiza citrinella) and pine bunting (Emberiza leucocephala) as an evidence of population structure and evolutionary history of the species, Zoologicheskii Zhurnal, 86(7), 863–876.

65. Rubtsov, A. S. (2022). Pair composition, habitat preferences and comparative life durations of birds in a hybrid population of the yellowhammer (Emberiza citrinella) and the pine bunting (E. leucocephalos) (Passeriformes, Emberizidae) in the Altais. Biology Bulletin, 49, (8), 1186–1196.

66. Rubtsov, A. S., & Tarasov, V. V. (2017). Relations between the yellowhammer (Emberiza citrinella) and the pine bunting (Emberiza leucocephalos) in the forested steppe of the Trans-Urals. Biology Bulletin, 44(9), 1059–1072. https://doi:10.1134/S1062359017090114

67. Saetre, G.P., Moum, T., Bureš, S., Král, M., Adamjan, M., & Moreno, J. (1997). A sexually selected character displacement in flycatchers reinforces premating isolation. Nature 387(6633), 589–592. https://doi:10.1038/42451

68. Sambrook, J., Fritsch, E.F., & Maniatis, T. (1989). Molecular cloning: a laboratory manual (No. Ed. 2). Cold spring harbor laboratory press.

69. Sarkar, D. (2008). Lattice: Multivariate data visualization with R. Springer. https://doi:10.1007/978-0-387-75969-2

70. Scheet, P., & Stephens, M. (2006). A fast and flexible statistical model for large-scale population genotype data: Applications to inferring missing genotypes and haplotypic phase. American Journal of Human Genetics, 78(4), 629–644. https://doi:10.1086/502802

71. Seehausen, O., & van Alphen, Jacques J. M. (1998). The effect of male coloration on female mate choice in closely related Lake Victoria cichlids (Haplochromis nyererei complex). Behavioral Ecology and Sociobiology, 42(1), 1–8. https://doi:10.1007/s002650050405

72. Southern, H. N. (1945). Polymorphism in Poephila gouldiae gould. Journal of Genetics, 47(1), 51–57.

73. Stacklies, W., Redestig, H., Scholz, M., Walther, D., & Selbig, J. (2007). pcaMethods-a bioconductor package providing PCA methods for incomplete data. Bioinformatics, 23(9), 1164–1167. https://doi:10.1093/bioinformatics/btm069

74. Strunnikova, N. V., Maminishkis, A., Barb, J. J., Wang, F., Zhi, C., Sergeev, Y., Chen, W., Edwards, A. O., Stambolian, D., Abecasis, G., Swaroop, A., Munson, P. J., & Miller, S. S. (2010). Transcriptome analysis and molecular signature of human retinal pigment epithelium. Human Molecular Genetics, 19(12), 2468–2486. https://doi:10.1093/hmg/ddq129

75. Theron, E., Hawkins, K., Bermingham, E., Ricklefs, R. E., & Mundy, N. I. (2001). The molecular basis of an avian plumage polymorphism in the wild: A melanocortin-1-receptor point mutation is perfectly associated with the melanic plumage morph of the bananaquit, Coereba flaveola. Current Biology, 11(8), 550–557. https://doi.org/10.1016/S0960-9822(01)00158-0

76. Todesco, M., Owens, G. L., Bercovich, N., Légaré, J., Soudi, S., Burge, D. O., Huang, K., Ostevik, K. L., Drummond, E. B. M., Imerovski, I., Lande, K., Pascual-Robles, M. A., Nanavati, M., Jahani, M., Cheung, W., Staton, S. E., Muños, S., Nielsen, R., Donovan, L. A., … Rieseberg, L. H. (2020). Massive haplotypes underlie ecotypic differentiation in sunflowers. Nature, 584(7822), 602–607. https://doi:10.1038/s41586-020-2467-6

77. Toews, D. L., Taylor, S., Vallender, R., Brelsford, A., Butcher, B., Messer, P., & Lovette, I. (2016). Plumage genes and little else distinguish the genomes of hybridizing warblers. Current Biology, 26(17), 2313–2318. https://doi:10.1016/j.cub.2016.06.034

78. Toomey, M. B., Lopes, R. J., Araújo, P. M., Johnson, J. D., Gazda, M. A., Afonso, S., Mota, P. G., Koch, R. E., Hill, G. E., Corbo, J. C., & Carneiro, M. (2017). High-density lipoprotein receptor SCARB1 is required for carotenoid coloration in birds. Proceedings of the National Academy of Sciences, 114(20), 5219–5224. https://doi:10.1073/pnas.1700751114

79. Turbek, S. P., Browne, M., Di Giacomo, A. S., Kopuchian, C., Hochachka, W. M., Estalles, C., Lijtmaer, D. A., Tubaro, P. L., Silveira, L. F., Lovette, I. J., Safran, R. J., Taylor, S. A., & Campagna, L. (2021). Rapid speciation via the evolution of pre-mating isolation in the iberá seedeater. Science, 371(6536), eabc0256. https://doi:10.1126/science.abc0256

80. Turner, S. (2018). qqman: An R package for visualizing GWAS results using Q-Q and Manhattan plots. Journal of Open Source Software, 3(25), 731. https://doi:10.21105/joss.00731

81. Tuttle, E. M. (2003). Alternative reproductive strategies in the white-throated sparrow: Behavioral and genetic evidence. Behavioral Ecology, 14(3), 425–432. https://doi:10.1093/beheco/14.3.425

82. Tuttle, E., Bergland, A., Korody, M., Brewer, M., Newhouse, D., Minx, P., Stager, M., Betuel, A., Cheviron, Z., Warren, W., Gonser, R., & Balakrishnan, C. (2016). Divergence and functional degradation of a sex chromosome-like supergene. Current Biology, 26(3), 344–350. https://doi:10.1016/j.cub.2015.11.069

83. Uy, J. A. C., Moyle, R. G., & Filardi, C. E. (2009). Plumage and song differences mediate species recognition between incipient flycatcher species of the Solomon Islands. Evolution, 63(1), 153–164. https://doi:10.1111/j.1558-5646.2008.00530.x

84. van der Sluijs, I., Van Dooren, T. J. M., Hofker, K. D., van Alphen, J. J. M., Stelkens, R. B., & Seehausen, O. (2008). Female mating preference functions predict sexual selection against hybrids between sibling species of cichlid fish. Philosophical Transactions: Biological Sciences, 363(1505), 2871–2877. https://doi:10.1098/rstb.2008.0045

85. Wang, I. J., & Shaffer, H. B. (2008). Rapid color evolution in an aposematic species: A phylogenetic analysis of color variation in the strikingly polymorphic strawberry poison-dart frog. Evolution, 62(11), 2742–2759. https://doi:10.1111/j.1558-5646.2008.00507.x

86. Warren, W. C., Clayton, D. F., Ellegren, H., Arnold, A. P., Hillier, L. W., Künstner, A., Searle, S., White, S., Vilella, A. J., Fairley, S., Heger, A., Kong, L., Ponting, C. P., Jarvis, E. D., Mello, C. V., Minx, P., Lovell, P., Velho, T. A. F., Ferris, M., … Wilson, R. K. (2010). The genome of a songbird. Nature, 464(7289), 757–762. https://doi:10.1038/nature08819

87. Weir, B. S., & Cockerham, C. C. (1984). Estimating F-statistics for the analysis of population structure. Evolution, 38(6), 1358–1370. https://doi:10.1111/j.1558-5646.1984.tb05657.x

88. West-Eberhard, M. J. (1983). Sexual selection, social competition, and speciation. The Quarterly Review of Biology, 58(2), 155–183. https://doi:10.1086/413215

89. Widelitz, R. B., Jiang, T., Lu, J., & Chuong, C. (2000). β-catenin in epithelial morphogenesis: Conversion of part of avian foot scales into feather buds with a mutated β-catenin. Developmental Biology, 219(1), 98–114. https://doi:10.1006/dbio.1999.9580

90. Xu, J., Lin, S., Gao, X., Nie, Q., Luo, Q., & Zhang, X. (2017). Mapping of id locus for dermal shank melanin in a Chinese indigenous chicken breed. Journal of Genetics, 96(6), 977–983. https://doi:10.1007/s12041-017-0862-z

91. Zheng, X., Levine, D., Shen, J., Gogarten, S. M., Laurie, C., & Weir, B. S. (2012). A high-performance computing toolset for relatedness and principal component analysis of SNP data. Bioinformatics, 28(24), 3326–3328. https://doi:10.1093/bioinformatics/bts606

92. Zhou, X., & Stephens, M. (2012). Genome-wide efficient mixed-model analysis for association studies. Nature Genetics, 44(7), 821–824. https://doi:10.1038/ng.2310

