## Supplementary Materials for "Evidence for a chromosomal inversion maintaining divergent plumage phenotypes between extensively hybridizing yellowhammers (*Emberiza citrinella*) and pine buntings (*E. leucocephalos*)"

### *Supplemental Text*

#### A discussion of PCA outliers

In a PCA of SNP information from individuals across the yellowhammer and pine bunting system (Supplementary Figure 3) as well as in PCAs that contain genetic information from subsets of individuals (Supplementary Figure 1: Supplementary Figure 2), we identified several outliers sometimes appearing as outlier pairs. Such a pattern could indicate kinship between outliers that would affect PCA results. To address this possibility, we conducted a kinship analysis of all analyzed samples and found that the majority of individuals possessed no kinship relationship (Supplementary Figure 4). Three pairs of individuals showed some amount of relatedness (Supplementary Table 2), but only one appeared as an outlier pair in the PCA depicted in Supplementary Figure 2. As such, kinship does not appear to have significantly affected PCA results and, most importantly, the trends observed in Figure 2A. It is unclear what forces may be driving these outlier patterns, but, in any case, outlier removal did not compromise system-wide genetic clustering patterns and merely created more noise along the secondary PC axis (Supplementary Figure 5).

***Supplemental Figures***

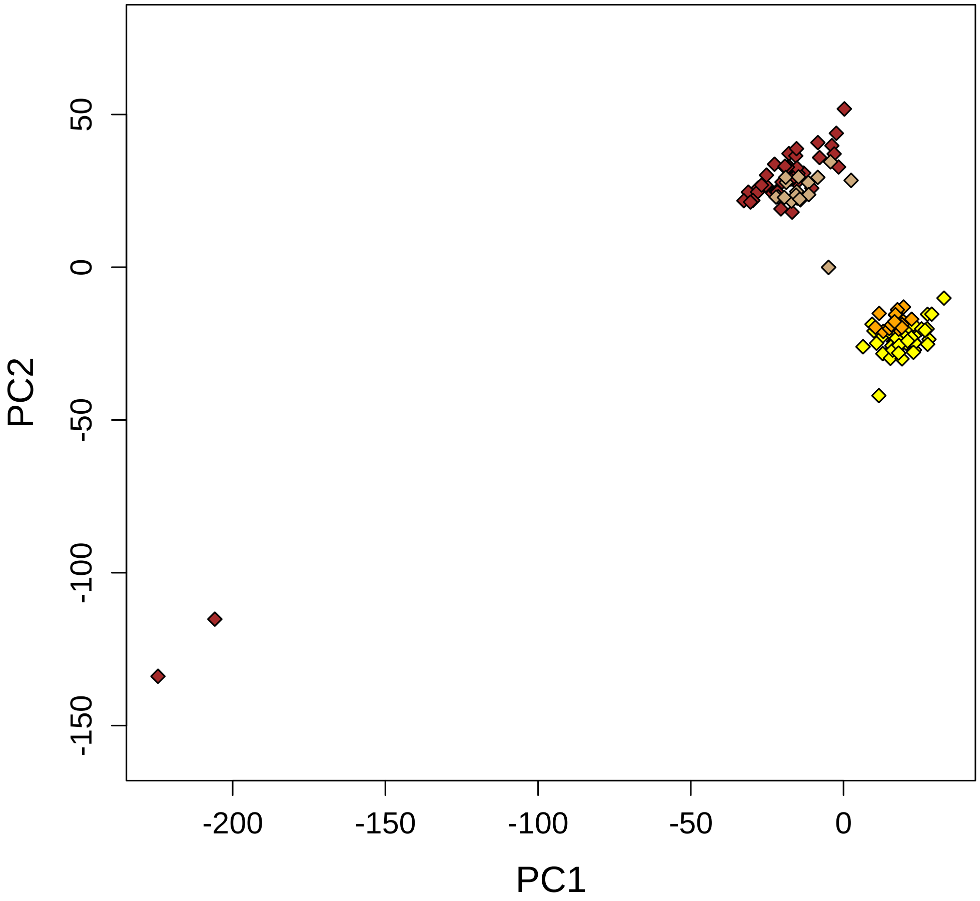


**Supplementary Figure 1.** Whole-genome principal components analysis of allopatric yellowhammers (yellow; n = 53), near-sympatric yellowhammers (light orange; n = 15), allopatric pine buntings (brown; n = 42) and near-sympatric pine buntings (taupe; n = 18). Colour legend available in Figure 2A. PC1 explains 2.9% of the variation among individuals and PC2 explains 2.5% of the variation among individuals. Information from 374,780 SNPs was included in this analysis.

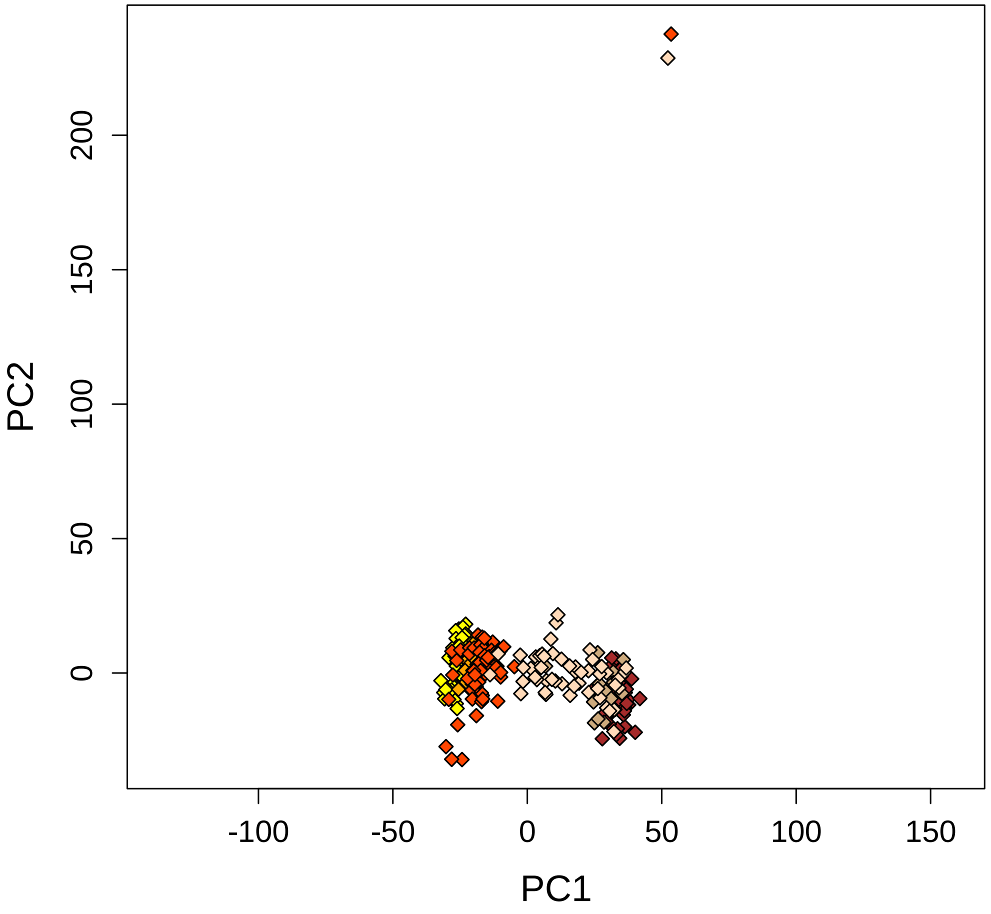


**Supplementary Figure 2.** Whole-genome principal components analysis of allopatric yellowhammers (yellow; n = 53), near-sympatric yellowhammers (light orange; n = 15), sympatric yellowhammers (red-orange; n = 67), allopatric pine buntings (brown; n = 42), near-sympatric pine buntings (taupe; n = 18) and sympatric pine buntings (peach; n = 52). Colour legend available in Figure 2A. PC1 explains 1.7% of the variation among individuals and PC2 explains 1.5% of the variation among individuals. Information from 374,780 SNPs was included in this analysis.

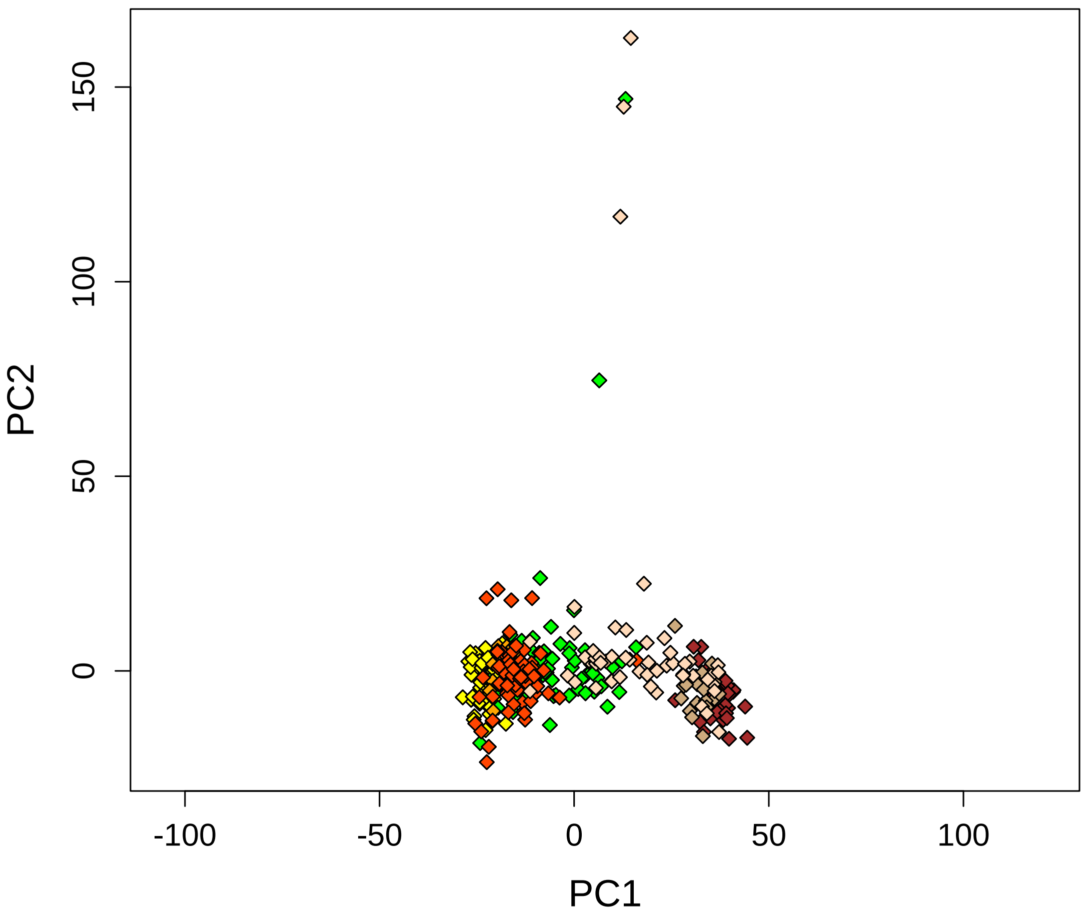


**Supplementary Figure 3.** Whole-genome principal components analysis of allopatric yellowhammers (yellow; n = 53), near-sympatric yellowhammers (light orange; n = 15), sympatric yellowhammers (red-orange; n = 67), allopatric pine buntings (brown; n = 42), near-sympatric pine buntings (taupe; n =18), sympatric pine buntings (peach; n = 52) and hybrids (green; n = 74). Colour legend available in Figure 2A. PC1 explains 1.4% of the variation among individuals and PC2 explains 0.9% of the variation among individuals. Information from 374,780 SNPs was included in this analysis.

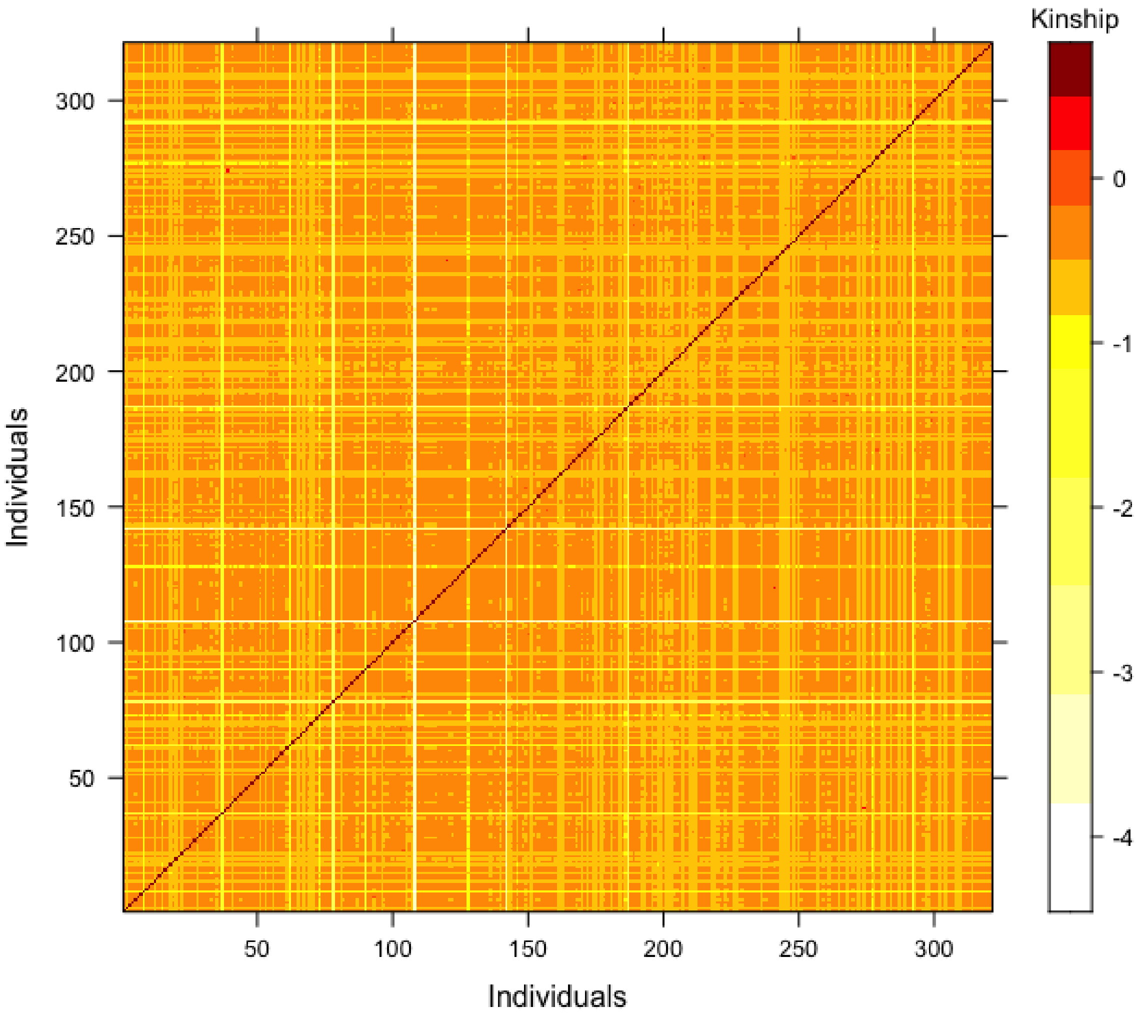


**Supplementary Figure 4.** Heatmap showing kinship relationships between all individuals included in this study. Kinship relationships were calculated as identity-by-descent coefficients using the KING method. High coefficients indicate a closer kinship relationship between individuals. Results on either side of the red diagonal are identical.

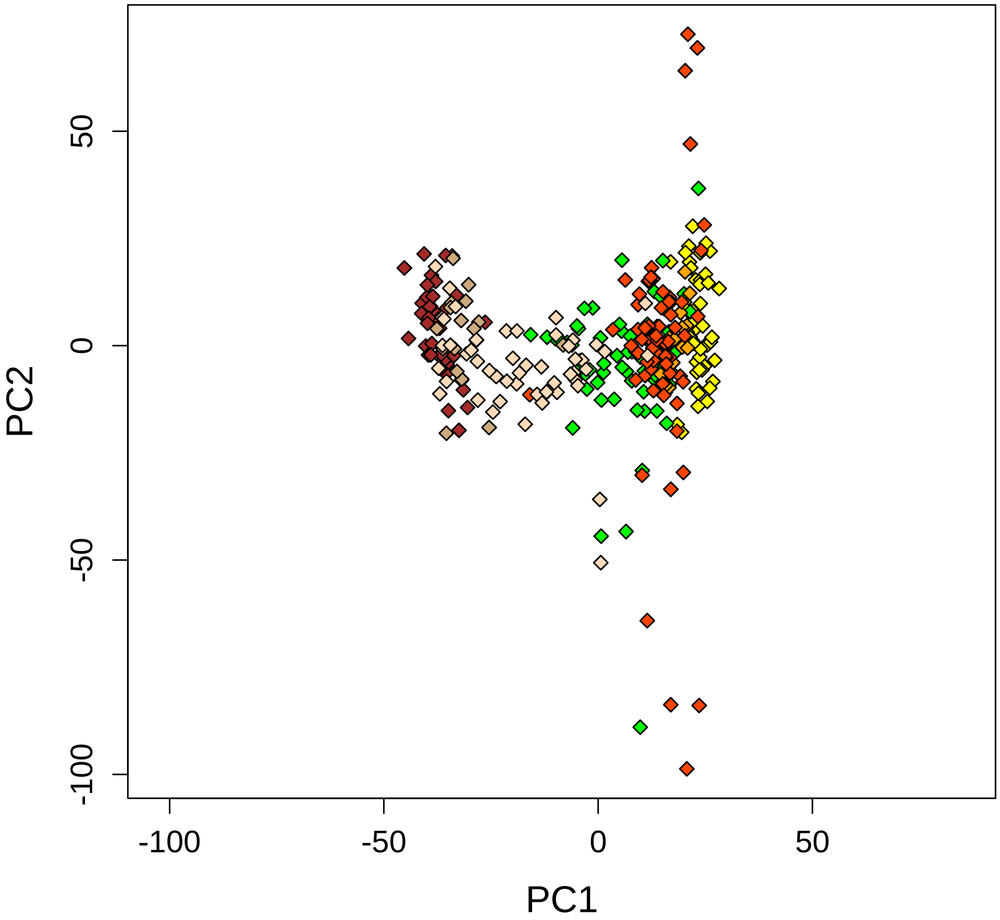


**Supplementary Figure 5.** Whole-genome principal components analysis of allopatric yellowhammers (yellow; n = 53), near-sympatric yellowhammers (light orange; n = 15), sympatric yellowhammers (red-orange; n = 67), allopatric pine buntings (brown; n = 42), near-sympatric pine buntings (taupe; n = 18), sympatric pine buntings (peach; n = 49) and hybrids (green; n = 72) following the removal of five outliers identified in Supplementary Figure 5: “Emberiza_GBS4_XD_632”, “Emberiza_GBS4_XD_636”, “Emberiza_GBS4_XD_639”, “Emberiza_GBS4_XD_972” and “Emberiza_GBS5_XD_970”. Colour legend available in Figure 2A. PC1 explains 1.4% of the variation among individuals and PC2 explains 0.8% of the variation among individuals. Information from 374,780 SNPs was included in this analysis.

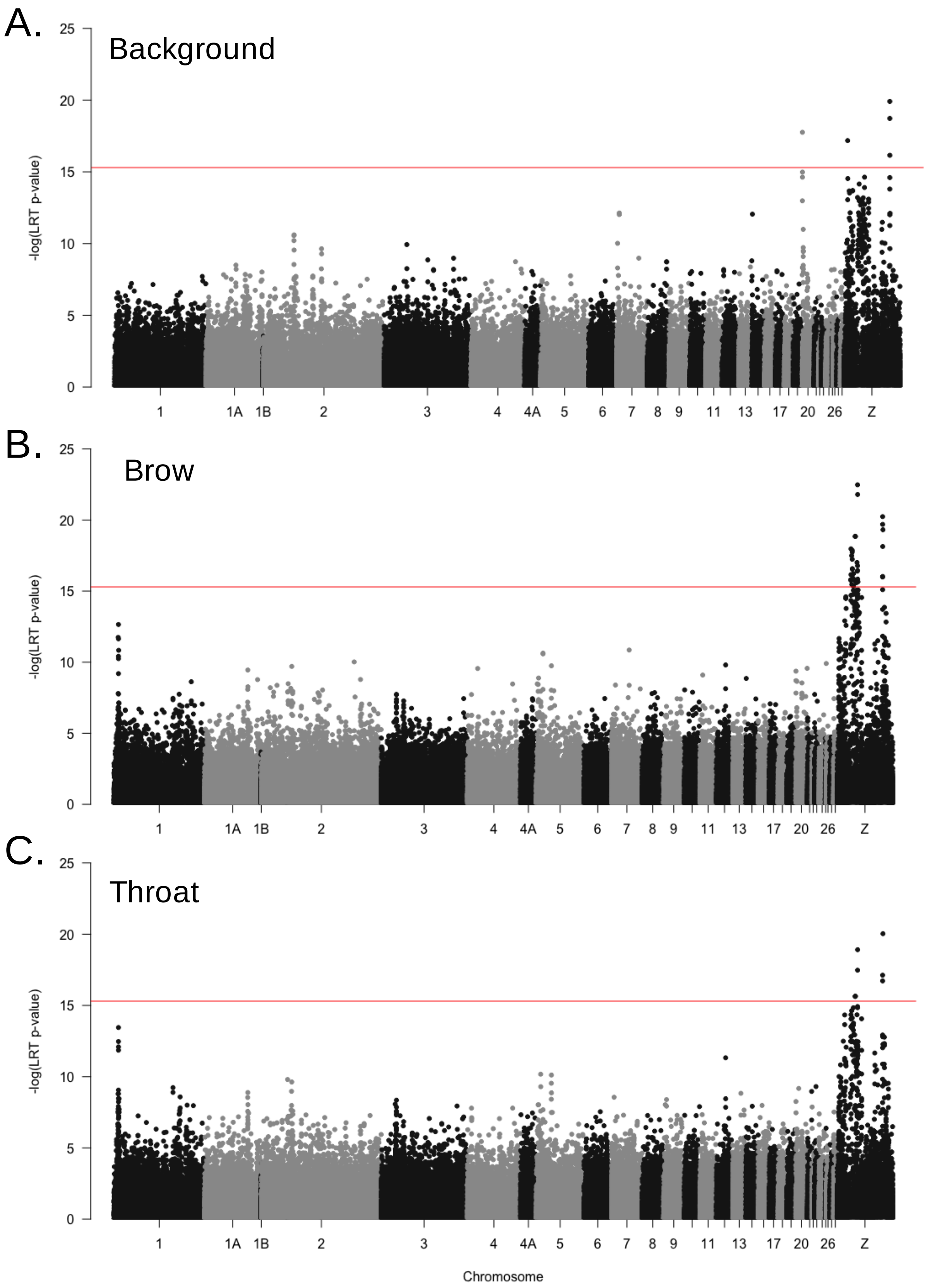
A.
Background
B.
Brow
C.
Throat
-log(LRT p-value)
0
5
10
15
20
25
1
1A
1B
2
3
4
4A
5
6
7
8
9
11
13
17
20
26
Z
Chromosome

**Supplementary Figure 6.** Associations between genome-wide SNPs and phenotypic variation in three plumage traits within the yellowhammer and pine bunting system. P-values for each SNP were determined using a likelihood ratio test calculated in the GEMMA program. Red lines indicate Bonferroni corrected significance thresholds. **A)** Associations between 220,220 genome-wide SNPs and variation in the background plumage colour. **B)** Associations between 220,124 genome-wide SNPs and variation in the amount of chestnut plumage at the brow. **C)** Associations between 220,307 genome-wide SNPs and variation in the amount of chestnut plumage at the throat.

*Supplemental Tables*

**Supplementary Table 1.** Detailed information on all the samples included in this study. Where possible, museum accession numbers were included as part of the sample IDs following “Emberiza_GBS#_”. Otherwise, samples were coded based on the museum they were obtained from or the collector’s coding system. Explanations for the abbreviations used in the “Pheno Class” (Phenotypic Class) and “Geographic Distribution” columns can be found in the Methods section. In the “sex” column, “m” stands for male, “f” stands for female and “uk” stands for unknown. The “TH” column contains phenotypic scores for each individual for the throat plumage trait. The “BR” column contains phenotypic scores for each individual for the brow plumage trait. The “BG” column contains phenotypic scores for each individual for the background colour plumage trait. In the “Pheno Class” column, “FML” stands for female and “UK” stands for unknown. The numbers in the “Sampling Location” column correspond to those that appear in Figure 1A. In all columns, a “NA” observation stands for “Not Applicable”.

| Sample ID | Species | Sex | TH | BR | BG | Pheno Class | Geographic Distribution | Source | Latitude (°N) | Longitude (°E) | Sampling Location |
|---|---|---|---|---|---|---|---|---|---|---|---|
| Emberiza_GBS1_ASR00_01 | E. leucocephalos | m | 7 | 7 | 7 | PL | Allopatric | Collected | 51.26 | 115.21 | 28 |
| Emberiza_GBS1_ASR05_14 | E. citrinella | m | 1 | 2 | 1 | SC | Allopatric | Collected | 51.2 | 57.27 | 12 |
| Emberiza_GBS1_ASR05_17 | E. citrinella | m | 0 | 0 | 0 | PC | Allopatric | Collected | 51.2 | 57.27 | 12 |
| Emberiza_GBS1_ASR05_18 | E. citrinella | m | 2 | 1 | 1 | SC | Allopatric | Collected | 51.2 | 57.27 | 12 |
| Emberiza_GBS1_ASR05_33 | E. leucocephalos | m | 7 | 7 | 7 | PL | Allopatric | Collected | 51.12 | 118.56 | 29 |
| Emberiza_GBS1_ASR05_35 | E. leucocephalos | m | 7 | 7 | 7 | PL | Allopatric | Collected | 51.12 | 118.56 | 29 |
| Emberiza_GBS1_ASR05_36 | E. leucocephalos | m | 7 | 7 | 7 | PL | Allopatric | Collected | 51.12 | 118.56 | 29 |
| Emberiza_GBS1_ASR05_37 | E. leucocephalos | m | 7 | 7 | 7 | PL | Allopatric | Collected | 51.12 | 118.56 | 29 |
| Emberiza_GBS1_ASR05_43 | E. leucocephalos | m | 7 | 7 | 7 | PL | Allopatric | Collected | 51.12 | 118.56 | 29 |
| Emberiza_GBS1_ASR05_45 | E. leucocephalos | m | 7 | 7 | 7 | PL | Allopatric | Collected | 51.12 | 118.56 | 29 |
| Emberiza_GBS1_ASR05_47 | E. leucocephalos | m | 7 | 7 | 7 | PL | Allopatric | Collected | 51.12 | 118.56 | 29 |
| Emberiza_GBS1_ASR05_54 | E. leucocephalos | m | 7 | 7 | 7 | PL | Allopatric | Collected | 50.21 | 115.06 | 28 |
| Emberiza_GBS1_ASR05_55 | E. leucocephalos | m | 7 | 7 | 7 | PL | Allopatric | Collected | 50.21 | 115.06 | 28 |
| Emberiza_GBS1_ASR05_56 | E. leucocephalos | m | 7 | 7 | 7 | PL | Allopatric | Collected | 50.21 | 115.06 | 28 |
| Emberiza_GBS1_ASR05_59 | E. leucocephalos | m | 7 | 7 | 7 | PL | Allopatric | Collected | 50.21 | 115.06 | 28 |

| Sample ID | Species | Sex | TH | BR | BG | Pheno Class | Geographic Distribution | Source | Latitude (°N) | Longitude (°E) | Sampling Location |
|---|---|---|---|---|---|---|---|---|---|---|---|
| Emberiza_GBS1_ASR05_61 | E. leucocephalos | m | 7 | 7 | 7 | PL | Allopatric | Collected | 50.21 | 115.06 | 28 |
| Emberiza_GBS1_ASR05_66 | E. leucocephalos | m | 7 | 7 | 7 | PL | Allopatric | Collected | 50.21 | 115.06 | 28 |
| Emberiza_GBS1_ASR05_68 | E. leucocephalos | m | 7 | 7 | 7 | PL | Allopatric | Collected | 50.21 | 115.06 | 28 |
| Emberiza_GBS1_AWH_067 | E. citrinella | f | NA | NA | NA | FML | Sympatric | Burke Museum, USA | 52.528 | 104.297 | 24 |
| Emberiza_GBS1_AWH_076 | E. leucocephalos | m | 7 | 7 | 7 | PL | Sympatric | Burke Museum, USA | 52.528 | 104.297 | 24 |
| Emberiza_GBS1_AWH_077 | E. leucocephalos | f | NA | NA | NA | FML | Sympatric | Burke Museum, USA | 52.528 | 104.297 | 24 |
| Emberiza_GBS1_AWH_148 | E. citrinella | f | NA | NA | NA | FML | Sympatric | Burke Museum, USA | 56.405 | 105.463 | 25 |
| Emberiza_GBS1_AWH_169 | E. citrinella | f | NA | NA | NA | FML | Sympatric | Burke Museum, USA | 56.405 | 105.463 | 25 |
| Emberiza_GBS1_BKS_1572 | E. citrinella | m | 1 | 0 | 0 | SC | Allopatric | Burke Museum, USA | 54.573 | 39.182 | 7 |
| Emberiza_GBS1_BKS_1646 | E. citrinella | f | NA | NA | NA | FML | Allopatric | Burke Museum, USA | 54.573 | 39.182 | 7 |
| Emberiza_GBS1_BKS_1821 | E. citrinella | m | 0 | 0 | 0 | PC | Allopatric | Burke Museum, USA | 51.412 | 34.547 | 6 |
| Emberiza_GBS1_BKS_1841 | E. citrinella | m | 1 | 1 | 1 | SC | Near Sympatric | Burke Museum, USA | 56.318 | 59.18 | 13 |

| Sample ID | Species | Sex | TH | BR | BG | Pheno Class | Geographic Distribution | Source | Latitude (°N) | Longitude (°E) | Sampling Location |
|---|---|---|---|---|---|---|---|---|---|---|---|
| Emberiza_GBS1_BKS_1928 | E. citrinella | m | 1 | 0 | 1 | SC | Near Sympatric | Burke Museum, USA | 56.456 | 58.065 | 13 |
| Emberiza_GBS1_BKS_1945 | E. citrinella | m | 1 | 0 | 0 | SC | Near Sympatric | Burke Museum, USA | 56.456 | 58.065 | 13 |
| Emberiza_GBS1_BKS_1957 | E. citrinella | m | 2 | 1 | 0 | SC | Near Sympatric | Burke Museum, USA | 56.456 | 58.065 | 13 |
| Emberiza_GBS1_BKS_1988 | E. citrinella | m | 0 | 0 | 0 | PC | Allopatric | Burke Museum, USA | 55.54 | 39.352 | 7 |
| Emberiza_GBS1_BKS_1989 | E. citrinella | m | 0 | 0 | 0 | PC | Allopatric | Burke Museum, USA | 55.54 | 39.352 | 7 |
| Emberiza_GBS1_BKS_2040 | E. citrinella | f | NA | NA | NA | FML | Allopatric | Burke Museum, USA | 55.54 | 39.352 | 7 |
| Emberiza_GBS1_BKS_2041 | E. citrinella | m | 1 | 0 | 0 | SC | Allopatric | Burke Museum, USA | 55.54 | 39.352 | 7 |
| Emberiza_GBS1_CDS_4883 | E. leucocephalos | uk | NA | NA | NA | UK | Near Sympatric | Burke Museum, USA | 51.346 | 106.5096 | 26 |
| Emberiza_GBS1_CDS_4906 | E. leucocephalos | f | NA | NA | NA | FML | Near Sympatric | Burke Museum, USA | 51.346 | 106.5096 | 26 |
| Emberiza_GBS1_DAB_296 | E. leucocephalos | m | 7 | 7 | 7 | PL | Near Sympatric | Burke Museum, USA | 51.346 | 106.5096 | 26 |
| Emberiza_GBS1_DAB_297 | E. leucocephalos | m | 7 | 7 | 7 | PL | Near Sympatric | Burke Museum, USA | 51.346 | 106.5096 | 26 |

| Sample ID | Species | Sex | TH | BR | BG | Pheno Class | Geographic Distribution | Source | Latitude (°N) | Longitude (°E) | Sampling Location |
|---|---|---|---|---|---|---|---|---|---|---|---|
| Emberiza_GBS1_ENP05_06 | E. citrinella | uk | NA | NA | NA | UK | Sympatric | Collected | 51.96 | 85.96 | 17 |
| Emberiza_GBS1_ENP05_08 | E. citrinella | uk | NA | NA | NA | UK | Sympatric | Collected | 51.96 | 85.96 | 17 |
| Emberiza_GBS1_ENP05_14 | E. citrinella | uk | NA | NA | NA | UK | Sympatric | Collected | 52.53 | 85.2 | 16 |
| Emberiza_GBS1_ENP05_16 | E. citrinella | uk | NA | NA | NA | UK | Sympatric | Collected | 53.35 | 83.75 | 16 |
| Emberiza_GBS1_ENP05_20 | E. citrinella | uk | NA | NA | NA | UK | Sympatric | Collected | 53.35 | 83.75 | 16 |
| Emberiza_GBS1_ENP05_22 | E. citrinella | uk | NA | NA | NA | UK | Sympatric | Collected | 53.35 | 83.75 | 16 |
| Emberiza_GBS1_ENP05_25 | E. leucocephalos | uk | NA | NA | NA | UK | Sympatric | Collected | 53.35 | 83.75 | 16 |
| Emberiza_GBS1_ENP97_11 | E. citrinella | uk | NA | NA | NA | UK | Sympatric | Collected | 54.85 | 83.11 | 16 |
| Emberiza_GBS1_ENP97_12 | Hybrid | uk | NA | NA | NA | UK | Sympatric | Collected | 54.85 | 83.11 | 16 |
| Emberiza_GBS1_ENP97_25 | E. citrinella | uk | NA | NA | NA | UK | Allopatric | Collected | 58.33 | 44.76 | 11 |
| Emberiza_GBS1_EVN_327 | E. citrinella | uk | NA | NA | NA | UK | Allopatric | Bell Museum, USA | 61.45 | 38.67 | 8 |
| Emberiza_GBS1_EVN_331 | E. citrinella | m | 0 | 0 | 2 | SC | Allopatric | Bell Museum, USA | 61.45 | 38.67 | 8 |
| Emberiza_GBS1_EVN_357 | E. citrinella | uk | NA | NA | NA | UK | Allopatric | Bell Museum, USA | 61.45 | 38.67 | 8 |
| Emberiza_GBS1_EVN_363 | E. citrinella | uk | NA | NA | NA | UK | Allopatric | Bell Museum, USA | 61.45 | 38.67 | 8 |
| Emberiza_GBS1_EVN_573 | E. citrinella | f | NA | NA | NA | FML | Allopatric | Darwin Museum, Russia | 57.71 | 39.34 | 7 |
| Emberiza_GBS1_EVN_574 | E. citrinella | m | 0 | 0 | 0 | PC | Allopatric | Darwin Museum, Russia | 57.71 | 39.34 | 7 |

| Sample ID | Species | Sex | TH | BR | BG | Pheno Class | Geographic Distribution | Source | Latitude (°N) | Longitude (°E) | Sampling Location |
|---|---|---|---|---|---|---|---|---|---|---|---|
| Emberiza_GBS1_IUK_615 | E. citrinella | m | 0 | 0 | 0 | PC | Allopatric | Bell Museum, USA | 61.45 | 38.67 | 8 |
| Emberiza_GBS1_IUK_631 | E. citrinella | f | NA | NA | NA | FML | Allopatric | Bell Museum, USA | 61.45 | 38.67 | 8 |
| Emberiza_GBS1_IVF_309 | Hybrid | m | 3 | 2 | 4 | WH | Allopatric | Bell Museum, USA | 61.45 | 38.67 | 8 |
| Emberiza_GBS1_IVF_390 | E. citrinella | uk | NA | NA | NA | UK | Allopatric | Bell Museum, USA | 61.45 | 38.67 | 8 |
| Emberiza_GBS1_IVF_682 | E. leucocephalos | m | 7 | 7 | 7 | PL | Allopatric | Drovetsky expedition | 50.5036 | 115.0029 | 28 |
| Emberiza_GBS1_JML_291 | E. leucocephalos | f | NA | NA | NA | FML | Sympatric | Burke Museum, USA | 52.528 | 104.297 | 24 |
| Emberiza_GBS1_M05_05 | E. citrinella | uk | NA | NA | NA | UK | Allopatric | Collected | 55.28 | 20.97 | 4 |
| Emberiza_GBS1_MSU_N247 | E. leucocephalos | uk | NA | NA | NA | UK | Near Sympatric | Burke Museum, USA | 50.38 | 95.1 | 22 |
| Emberiza_GBS1_MSU_N339 | E. leucocephalos | uk | NA | NA | NA | UK | Near Sympatric | Burke Museum, USA | 50.4343 | 91.4182 | 20 |
| Emberiza_GBS1_MSU_P66 | E. leucocephalos | uk | NA | NA | NA | UK | Near Sympatric | Burke Museum, USA | 50.13 | 95.09 | 22 |
| Emberiza_GBS1_NVN_015 | E. citrinella | m | 0 | 0 | 0 | PC | Allopatric | Darwin Museum, Russia | 57.71 | 39.34 | 7 |

| Sample ID | Species | Sex | TH | BR | BG | Pheno Class | Geographic Distribution | Source | Latitude (°N) | Longitude (°E) | Sampling Location |
|---|---|---|---|---|---|---|---|---|---|---|---|
| Emberiza_GBS1_RCF_1807 | E. leucocephalos | m | 7 | 7 | 7 | PL | Near Sympatric | Burke Museum, USA | 50.39 | 91.36 | 20 |
| Emberiza_GBS1_RCF_1971 | E. leucocephalos | uk | NA | NA | NA | UK | Near Sympatric | Burke Museum, USA | 50.04 | 95.08 | 22 |
| Emberiza_GBS1_RCF_2235 | E. citrinella | m | 1 | 0 | 0 | SC | Sympatric | Burke Museum, USA | 52.528 | 104.297 | 24 |
| Emberiza_GBS1_SVD_2134 | E. citrinella | m | 0 | 0 | 0 | PC | Allopatric | Burke Museum, USA | 43.54 | 40.47 | 9 |
| Emberiza_GBS1_SVD_2695 | E. citrinella | uk | NA | NA | NA | UK | Allopatric | Bell Museum, USA | 61.45 | 38.67 | 8 |
| Emberiza_GBS1_SVD_3479 | E. leucocephalos | m | 7 | 7 | 7 | PL | Allopatric | Drovetsky expedition | 49.6439 | 110.1652 | 27 |
| Emberiza_GBS1_SVD_518 | E. leucocephalos | m | 7 | 7 | 7 | PL | Sympatric | Burke Museum, USA | 52.46 | 104.41 | 24 |
| Emberiza_GBS1_SVD_519 | E. leucocephalos | f | NA | NA | NA | FML | Sympatric | Burke Museum, USA | 52.46 | 104.41 | 24 |
| Emberiza_GBS1_SVD_531 | E. leucocephalos | m | 7 | 7 | 7 | PL | Sympatric | Burke Museum, USA | 52.46 | 104.41 | 24 |
| Emberiza_GBS1_SWM_03 | E. citrinella | uk | NA | NA | NA | UK | Allopatric | Swedish NHM | 57.99 | 12.49 | 1 |
| Emberiza_GBS1_SWM_10 | E. citrinella | uk | NA | NA | NA | UK | Allopatric | Swedish NHM | 65.86 | 21.48 | 5 |
| Emberiza_GBS1_SWM_12 | E. citrinella | uk | NA | NA | NA | UK | Allopatric | Swedish NHM | 65.86 | 21.48 | 5 |

| Sample ID | Species | Sex | TH | BR | BG | Pheno Class | Geographic Distribution | Source | Latitude (°N) | Longitude (°E) | Sampling Location |
|---|---|---|---|---|---|---|---|---|---|---|---|
| Emberiza_GBS1_VGR_400 | E. citrinella | m | 1 | 0 | 0 | SC | Sympatric | Burke Museum, USA | 52.528 | 104.297 | 24 |
| Emberiza_GBS1_VGR_440 | E. citrinella | m | 1 | 0 | 1 | SC | Sympatric | Burke Museum, USA | 56.405 | 105.463 | 25 |
| Emberiza_GBS1_VGR_508 | E. citrinella | m | 0 | 0 | 1 | SC | Sympatric | Burke Museum, USA | 56.405 | 105.463 | 25 |
| Emberiza_GBS1_VM_282a | E. leucocephalos | m | 7 | 7 | 7 | PL | Allopatric | Burke Museum, USA | 50.44 | 143.18 | 30 |
| Emberiza_GBS2_ASR05_02 | E. citrinella | m | 1 | 0 | 0 | SC | Allopatric | Collected | 51.2 | 57.27 | 12 |
| Emberiza_GBS2_ASR05_15 | E. citrinella | m | 0 | 0 | 0 | PC | Allopatric | Collected | 51.2 | 57.27 | 12 |
| Emberiza_GBS2_ASR05_20 | E. citrinella | m | 2 | 1 | 0 | SC | Allopatric | Collected | 51.2 | 57.27 | 12 |
| Emberiza_GBS2_ASR05_22 | E. citrinella | m | 2 | 1 | 0 | SC | Allopatric | Collected | 51.2 | 57.27 | 12 |
| Emberiza_GBS2_ASR05_32 | E. leucocephalos | m | 7 | 7 | 7 | PL | Allopatric | Collected | 51.12 | 118.56 | 29 |
| Emberiza_GBS2_ASR05_38 | E. leucocephalos | m | 7 | 7 | 7 | PL | Allopatric | Collected | 51.12 | 118.56 | 29 |
| Emberiza_GBS2_ASR05_39 | E. leucocephalos | m | 7 | 7 | 7 | PL | Allopatric | Collected | 51.12 | 118.56 | 29 |
| Emberiza_GBS2_ASR05_40 | E. leucocephalos | m | 7 | 7 | 7 | PL | Allopatric | Collected | 51.12 | 118.56 | 29 |
| Emberiza_GBS2_ASR05_41 | E. leucocephalos | m | 7 | 7 | 7 | PL | Allopatric | Collected | 51.12 | 118.56 | 29 |
| Emberiza_GBS2_ASR05_44 | E. leucocephalos | m | 7 | 7 | 7 | PL | Allopatric | Collected | 51.12 | 118.56 | 29 |
| Emberiza_GBS2_ASR05_46 | E. leucocephalos | f | NA | NA | NA | FML | Allopatric | Collected | 51.12 | 118.56 | 29 |
| Emberiza_GBS2_ASR05_49 | E. leucocephalos | m | 7 | 7 | 7 | PL | Allopatric | Collected | 51.12 | 118.56 | 29 |
| Emberiza_GBS2_ASR05_51 | E. leucocephalos | m | 7 | 7 | 7 | PL | Allopatric | Collected | 50.21 | 115.06 | 28 |
| Emberiza_GBS2_ASR05_52 | E. leucocephalos | m | 7 | 7 | 7 | PL | Allopatric | Collected | 50.21 | 115.06 | 28 |
| Emberiza_GBS2_ASR05_57 | E. leucocephalos | m | 7 | 7 | 7 | PL | Allopatric | Collected | 50.21 | 115.06 | 28 |

| Sample ID | Species | Sex | TH | BR | BG | Pheno Class | Geographic Distribution | Source | Latitude (°N) | Longitude (°E) | Sampling Location |
|---|---|---|---|---|---|---|---|---|---|---|---|
| Emberiza_GBS2_ASR05_58 | E. leucocephalos | m | 7 | 7 | 7 | PL | Allopatric | Collected | 50.21 | 115.06 | 28 |
| Emberiza_GBS2_ASR05_60 | E. leucocephalos | m | 7 | 7 | 7 | PL | Allopatric | Collected | 50.21 | 115.06 | 28 |
| Emberiza_GBS2_ASR05_67 | E. leucocephalos | m | 7 | 7 | 7 | PL | Allopatric | Collected | 50.21 | 115.06 | 28 |
| Emberiza_GBS2_ASR98_11 | Hybrid | m | 4 | 2 | 0 | CH | Sympatric | Darwin Museum, Russia | 53.32 | 107.12 | 26 |
| Emberiza_GBS2_ASR98_17 | E. leucocephalos | m | 7 | 7 | 7 | PL | Sympatric | Darwin Museum, Russia | 52.8 | 104.74 | 24 |
| Emberiza_GBS2_ASR99_01 | Hybrid | m | 0 | 0 | 4 | WH | Sympatric | Darwin Museum, Russia | 55.33 | 93.65 | 21 |
| Emberiza_GBS2_AWH_039 | E. citrinella | f | NA | NA | NA | FML | Sympatric | Burke Museum, USA | 52.528 | 104.297 | 24 |
| Emberiza_GBS2_AWH_152 | E. citrinella | f | NA | NA | NA | FML | Sympatric | Burke Museum, USA | 56.405 | 105.463 | 25 |
| Emberiza_GBS2_BKS_1583 | E. citrinella | m | 1 | 1 | 0 | SC | Allopatric | Burke Museum, USA | 54.573 | 39.182 | 7 |
| Emberiza_GBS2_BKS_1609 | E. citrinella | m | 1 | 0 | 0 | SC | Allopatric | Burke Museum, USA | 54.573 | 39.182 | 7 |
| Emberiza_GBS2_BKS_1654 | E. citrinella | m | 2 | 1 | 0 | SC | Allopatric | Burke Museum, USA | 51.349 | 37.123 | 6 |
| Emberiza_GBS2_BKS_1667 | E. citrinella | f | NA | NA | NA | FML | Allopatric | Burke Museum, USA | 51.349 | 37.123 | 6 |

| Sample ID | Species | Sex | TH | BR | BG | Pheno Class | Geographic Distribution | Source | Latitude (°N) | Longitude (°E) | Sampling Location |
|---|---|---|---|---|---|---|---|---|---|---|---|
| Emberiza_GBS2_BKS_1710 | Hybrid | m | 4 | 3 | 1 | CH | Allopatric | Burke Museum, USA | 51.349 | 37.123 | 6 |
| Emberiza_GBS2_BKS_1859 | E. citrinella | m | 1 | 0 | 1 | SC | Near Sympatric | Burke Museum, USA | 56.318 | 59.18 | 13 |
| Emberiza_GBS2_BKS_2017 | E. citrinella | m | 1 | 0 | 1 | SC | Allopatric | Burke Museum, USA | 55.54 | 39.352 | 7 |
| Emberiza_GBS2_DAB_291 | E. leucocephalos | f | NA | NA | NA | FML | Near Sympatric | Burke Museum, USA | 51.346 | 106.5096 | 26 |
| Emberiza_GBS2_DAB_299 | E. leucocephalos | m | 7 | 7 | 7 | PL | Near Sympatric | Burke Museum, USA | 51.346 | 106.5096 | 26 |
| Emberiza_GBS2_DAB_301 | E. leucocephalos | m | 7 | 7 | 7 | PL | Near Sympatric | Burke Museum, USA | 51.346 | 106.5096 | 26 |
| Emberiza_GBS2_DAB_303 | E. leucocephalos | uk | NA | NA | NA | UK | Near Sympatric | Burke Museum, USA | 51.346 | 106.5096 | 26 |
| Emberiza_GBS2_DAB_308 | E. leucocephalos | f | NA | NA | NA | FML | Near Sympatric | Burke Museum, USA | 51.346 | 106.5096 | 26 |
| Emberiza_GBS2_ENP05_04 | E. citrinella | uk | NA | NA | NA | UK | Sympatric | Collected | 51.96 | 85.96 | 17 |
| Emberiza_GBS2_ENP05_09 | E. citrinella | uk | NA | NA | NA | UK | Sympatric | Collected | 51.96 | 85.96 | 17 |
| Emberiza_GBS2_ENP05_11 | E. citrinella | uk | NA | NA | NA | UK | Sympatric | Collected | 51.96 | 85.96 | 17 |
| Emberiza_GBS2_ENP05_12 | E. leucocephalos | uk | NA | NA | NA | UK | Sympatric | Collected | 51.96 | 85.96 | 17 |
| Emberiza_GBS2_ENP05_15 | E. citrinella | uk | NA | NA | NA | UK | Sympatric | Collected | 53.35 | 83.75 | 16 |
| Emberiza_GBS2_ENP05_21 | E. leucocephalos | uk | NA | NA | NA | UK | Sympatric | Collected | 53.35 | 83.75 | 16 |

| Sample ID | Species | Sex | TH | BR | BG | Pheno Class | Geographic Distribution | Source | Latitude (°N) | Longitude (°E) | Sampling Location |
|---|---|---|---|---|---|---|---|---|---|---|---|
| Emberiza_GBS2_ENP05_24 | E. leucocephalos | uk | NA | NA | NA | UK | Sympatric | Collected | 53.35 | 83.75 | 16 |
| Emberiza_GBS2_ENP05_26 | E. citrinella | uk | NA | NA | NA | UK | Sympatric | Collected | 53.35 | 83.75 | 16 |
| Emberiza_GBS2_ENP97_01 | E. leucocephalos | uk | NA | NA | NA | UK | Sympatric | Collected | 54.83 | 77.67 | 15 |
| Emberiza_GBS2_ENP97_03 | E. leucocephalos | uk | NA | NA | NA | UK | Sympatric | Collected | 54.83 | 77.67 | 15 |
| Emberiza_GBS2_ENP97_05 | E. citrinella | uk | NA | NA | NA | UK | Sympatric | Collected | 54.85 | 83.11 | 16 |
| Emberiza_GBS2_ENP97_07 | E. citrinella | uk | NA | NA | NA | UK | Sympatric | Collected | 54.85 | 83.11 | 16 |
| Emberiza_GBS2_ENP97_09 | E. citrinella | uk | NA | NA | NA | UK | Sympatric | Collected | 54.85 | 83.11 | 16 |
| Emberiza_GBS2_ENP97_10 | E. citrinella | uk | NA | NA | NA | UK | Sympatric | Collected | 54.85 | 83.11 | 16 |
| Emberiza_GBS2_ENP97_13 | E. citrinella | uk | NA | NA | NA | UK | Sympatric | Collected | 54.85 | 83.11 | 16 |
| Emberiza_GBS2_ENP97_15 | E. citrinella | uk | NA | NA | NA | UK | Sympatric | Collected | 54.85 | 83.11 | 16 |
| Emberiza_GBS2_ENP97_20 | Hybrid | uk | NA | NA | NA | UK | Sympatric | Collected | 54.85 | 83.11 | 16 |
| Emberiza_GBS2_ENP97_21 | Hybrid | uk | NA | NA | NA | UK | Sympatric | Collected | 54.85 | 83.11 | 16 |
| Emberiza_GBS2_EVN_328 | E. citrinella | uk | NA | NA | NA | UK | Allopatric | Bell Museum, USA | 61.45 | 38.67 | 8 |
| Emberiza_GBS2_EVN_366 | E. citrinella | uk | NA | NA | NA | UK | Allopatric | Bell Museum, USA | 61.45 | 38.67 | 8 |
| Emberiza_GBS2_EVN_570 | E. citrinella | m | 0 | 0 | 0 | PC | Allopatric | Darwin Museum, Russia | 57.71 | 39.34 | 7 |
| Emberiza_GBS2_IUK_2343 | E. leucocephalos | m | 7 | 7 | 7 | PL | Allopatric | Drovetsky expedition | 49.6439 | 110.1652 | 27 |
| Emberiza_GBS2_IUK_702 | E. citrinella | m | 0 | 0 | 0 | PC | Allopatric | Bell Museum, USA | 61.45 | 38.67 | 8 |

| Sample ID | Species | Sex | TH | BR | BG | Pheno Class | Geographic Distribution | Source | Latitude (°N) | Longitude (°E) | Sampling Location |
|---|---|---|---|---|---|---|---|---|---|---|---|
| Emberiza_GBS2_IUK_703 | E. citrinella | uk | NA | NA | NA | UK | Allopatric | Bell Museum, USA | 61.45 | 38.67 | 8 |
| Emberiza_GBS2_IUK_801 | E. citrinella | f | NA | NA | NA | FML | Allopatric | Bell Museum, USA | 65.85 | 44.24 | 10 |
| Emberiza_GBS2_M05_01 | E. citrinella | uk | NA | NA | NA | UK | Allopatric | Collected | 55.28 | 20.97 | 4 |
| Emberiza_GBS2_M05_03 | E. citrinella | uk | NA | NA | NA | UK | Allopatric | Collected | 55.28 | 20.97 | 4 |
| Emberiza_GBS2_M05_08 | E. citrinella | uk | NA | NA | NA | UK | Allopatric | Collected | 55.28 | 20.97 | 4 |
| Emberiza_GBS2_M05_10 | E. citrinella | uk | NA | NA | NA | UK | Allopatric | Collected | 55.28 | 20.97 | 4 |
| Emberiza_GBS2_MSU_N71 | E. leucocephalos | m | 7 | 7 | 7 | PL | Near Sympatric | Burke Museum, USA | 50.13 | 95.09 | 22 |
| Emberiza_GBS2_RCF_1949 | E. leucocephalos | m | 7 | 7 | 7 | PL | Near Sympatric | Burke Museum, USA | 50.44 | 90.01 | 20 |
| Emberiza_GBS2_RCF_1970 | E. leucocephalos | f | NA | NA | NA | FML | Near Sympatric | Burke Museum, USA | 50.04 | 95.08 | 22 |
| Emberiza_GBS2_RCF_2191b | E. citrinella | f | NA | NA | NA | FML | Sympatric | Burke Museum, USA | 52.528 | 104.297 | 24 |
| Emberiza_GBS2_RCF_2192 | E. citrinella | f | NA | NA | NA | FML | Sympatric | Burke Museum, USA | 52.528 | 104.297 | 24 |
| Emberiza_GBS2_RCF_2230 | E. leucocephalos | m | 6 | 7 | 7 | SL | Sympatric | Burke Museum, USA | 52.528 | 104.297 | 24 |
| Emberiza_GBS2_SVD_3563 | E. leucocephalos | m | 7 | 7 | 7 | PL | Allopatric | Drovetsky expedition | 50.5036 | 115.0029 | 28 |

| Sample ID | Species | Sex | TH | BR | BG | Pheno Class | Geographic Distribution | Source | Latitude (°N) | Longitude (°E) | Sampling Location |
|---|---|---|---|---|---|---|---|---|---|---|---|
| Emberiza_GBS2_SVD_514 | E. citrinella | m | 1 | 0 | 0 | SC | Sympatric | Burke Museum, USA | 52.46 | 104.41 | 24 |
| Emberiza_GBS2_SVD_524 | E. citrinella | uk | NA | NA | NA | UK | Sympatric | Burke Museum, USA | 52.46 | 104.41 | 24 |
| Emberiza_GBS2_SVD_533 | E. citrinella | uk | NA | NA | NA | UK | Sympatric | Burke Museum, USA | 57.28 | 97.18 | 23 |
| Emberiza_GBS2_SWM_11 | E. citrinella | uk | NA | NA | NA | UK | Allopatric | Swedish NHM | 59.81 | 17.05 | 2 |
| Emberiza_GBS2_VGR_343 | Hybrid | m | 3 | 2 | 0 | CH | Sympatric | Burke Museum, USA | 52.528 | 104.297 | 24 |
| Emberiza_GBS2_VGR_350 | E. leucocephalos | uk | NA | NA | NA | UK | Sympatric | Burke Museum, USA | 52.528 | 104.297 | 24 |
| Emberiza_GBS2_VGR_355 | E. leucocephalos | m | 6 | 7 | 7 | SL | Sympatric | Burke Museum, USA | 52.528 | 104.297 | 24 |
| Emberiza_GBS2_VM_285a | E. leucocephalos | m | 7 | 7 | 7 | PL | Allopatric | Burke Museum, USA | 50.44 | 143.18 | 30 |
| Emberiza_GBS2_ZMUC_09 | E. citrinella | uk | NA | NA | NA | UK | Allopatric | ZMUC, Denmark | 51.71 | 18.61 | 3 |
| Emberiza_GBS4_MIM_165 | E. leucocephalos | f | NA | NA | NA | FML | Allopatric | Zoological museum, Russia | 50.68 | 142.97 | 30 |
| Emberiza_GBS4_R06_01 | Hybrid | m | 3 | 1 | 0 | CH | Sympatric | Collected | 51.57 | 85.56 | 17 |
| Emberiza_GBS4_RYA_2397 | Hybrid | m | 2 | 0 | 7 | WH | Sympatric | Zoological museum, Russia | 53.48 | 78.82 | 15 |

| Sample ID | Species | Sex | TH | BR | BG | Pheno Class | Geographic Distribution | Source | Latitude (°N) | Longitude (°E) | Sampling Location |
|---|---|---|---|---|---|---|---|---|---|---|---|
| Emberiza_GBS4_RYA_2680 | E. leucocephalos | f | NA | NA | NA | FML | Allopatric | Zoological museum, Russia | 50.68 | 142.97 | 30 |
| Emberiza_GBS4_RYA_3178 | E. leucocephalos | m | NA | NA | NA | UK | Allopatric | Zoological museum, Russia | 50.68 | 142.97 | 30 |
| Emberiza_GBS4_SVN_2335 | E. leucocephalos | m | NA | NA | NA | UK | Allopatric | Zoological museum, Russia | 50.68 | 142.97 | 30 |
| Emberiza_GBS4_XD_548 | E. leucocephalos | m | 7 | 7 | 7 | PL | Sympatric | Collected | 50.73 | 86.34 | 17 |
| Emberiza_GBS4_XD_552 | Hybrid | f | NA | NA | NA | FML | Sympatric | Collected | 50.73 | 86.34 | 17 |
| Emberiza_GBS4_XD_603 | E. citrinella | m | 0 | 0 | 0 | PC | Sympatric | Collected | 50.73 | 86.34 | 17 |
| Emberiza_GBS4_XD_610 | E. citrinella | m | 0 | 0 | 0 | PC | Sympatric | Collected | 50.73 | 86.34 | 17 |
| Emberiza_GBS4_XD_612 | Hybrid | m | 2 | 0 | 4 | WH | Near Sympatric | Collected | 55.22 | 65.42 | 14 |
| Emberiza_GBS4_XD_613 | Hybrid | m | 4 | 4 | 6 | LH | Near Sympatric | Collected | 55.22 | 65.42 | 14 |
| Emberiza_GBS4_XD_617 | Hybrid | m | 5 | 5 | 7 | LH | Near Sympatric | Collected | 55.22 | 65.42 | 14 |
| Emberiza_GBS4_XD_619 | E. citrinella | m | 1 | 0 | 0 | SC | Near Sympatric | Collected | 55.22 | 65.42 | 14 |
| Emberiza_GBS4_XD_621 | E. citrinella | m | 0 | 0 | 0 | PC | Near Sympatric | Collected | 55.19 | 65.26 | 14 |
| Emberiza_GBS4_XD_623 | E. citrinella | m | 1 | 0 | 0 | SC | Near Sympatric | Collected | 55.19 | 65.26 | 14 |
| Emberiza_GBS4_XD_624 | E. citrinella | m | 0 | 0 | 0 | PC | Near Sympatric | Collected | 55.19 | 65.26 | 14 |
| Emberiza_GBS4_XD_625 | E. citrinella | m | 1 | 0 | 0 | SC | Near Sympatric | Collected | 55.19 | 65.26 | 14 |

| Sample ID | Species | Sex | TH | BR | BG | Pheno Class | Geographic Distribution | Source | Latitude (°N) | Longitude (°E) | Sampling Location |
|---|---|---|---|---|---|---|---|---|---|---|---|
| Emberiza_GBS4_XD_630 | Hybrid | m | 5 | 5 | 6 | LH | Near Sympatric | Collected | 54.67 | 64.88 | 14 |
| Emberiza_GBS4_XD_631 | E. citrinella | m | 1 | 0 | 0 | SC | Near Sympatric | Collected | 54.67 | 64.88 | 14 |
| Emberiza_GBS4_XD_632 | Hybrid | m | 4 | 3 | 2 | CH | Sympatric | Collected | 50.73 | 86.34 | 17 |
| Emberiza_GBS4_XD_634 | E. citrinella | m | 2 | 0 | 0 | SC | Sympatric | Collected | 50.73 | 86.34 | 17 |
| Emberiza_GBS4_XD_635 | E. leucocephalos | f | NA | NA | NA | FML | Sympatric | Collected | 50.73 | 86.34 | 17 |
| Emberiza_GBS4_XD_636 | E. leucocephalos | m | 7 | 7 | 7 | PL | Sympatric | Collected | 50.73 | 86.34 | 17 |
| Emberiza_GBS4_XD_637 | E. leucocephalos | m | 7 | 7 | 7 | PL | Sympatric | Collected | 50.73 | 86.34 | 17 |
| Emberiza_GBS4_XD_639 | Hybrid | m | 5 | 5 | 7 | LH | Sympatric | Collected | 50.73 | 86.34 | 17 |
| Emberiza_GBS4_XD_640 | E. leucocephalos | m | 7 | 7 | 4 | SL | Sympatric | Collected | 50.73 | 86.34 | 17 |
| Emberiza_GBS4_XD_645 | E. leucocephalos | m | 7 | 7 | 7 | PL | Sympatric | Collected | 50.73 | 86.34 | 17 |
| Emberiza_GBS4_XD_647 | E. citrinella | m | 0 | 0 | 0 | PC | Sympatric | Collected | 50.73 | 86.34 | 17 |
| Emberiza_GBS4_XD_649 | E. leucocephalos | m | 7 | 7 | 7 | PL | Sympatric | Collected | 50.73 | 86.34 | 17 |
| Emberiza_GBS4_XD_650 | Hybrid | m | 5 | 5 | 6 | LH | Sympatric | Collected | 50.73 | 86.34 | 17 |
| Emberiza_GBS4_XD_653 | E. leucocephalos | m | 7 | 7 | 7 | PL | Sympatric | Collected | 50.73 | 86.34 | 17 |
| Emberiza_GBS4_XD_654 | E. leucocephalos | m | 7 | 7 | 6 | SL | Sympatric | Collected | 50.73 | 86.34 | 17 |
| Emberiza_GBS4_XD_655 | Hybrid | m | 7 | 7 | 0 | YH | Sympatric | Collected | 50.73 | 86.34 | 17 |
| Emberiza_GBS4_XD_656 | Hybrid | m | 6 | 7 | 0 | YH | Sympatric | Collected | 50.73 | 86.34 | 17 |
| Emberiza_GBS4_XD_657 | E. citrinella | m | 1 | 0 | 0 | SC | Allopatric | Collected | 56.06 | 36.13 | 7 |
| Emberiza_GBS4_XD_658 | E. citrinella | m | 0 | 0 | 1 | SC | Allopatric | Collected | 56.06 | 36.13 | 7 |
| Emberiza_GBS4_XD_659 | E. citrinella | m | 1 | 1 | 0 | SC | Allopatric | Collected | 56.06 | 36.13 | 7 |
| Emberiza_GBS4_XD_663 | Hybrid | m | 1 | 2 | 5 | WH | Sympatric | Collected | 50.73 | 86.34 | 17 |
| Emberiza_GBS4_XD_666 | E. leucocephalos | m | 7 | 7 | 7 | PL | Sympatric | Collected | 50.73 | 86.34 | 17 |
| Emberiza_GBS4_XD_670 | E. citrinella | m | 0 | 0 | 2 | SC | Sympatric | Collected | 50.73 | 86.34 | 17 |

| Sample ID | Species | Sex | TH | BR | BG | Pheno Class | Geographic Distribution | Source | Latitude (°N) | Longitude (°E) | Sampling Location |
|---|---|---|---|---|---|---|---|---|---|---|---|
| Emberiza_GBS4_XD_672 | E. leucocephalos | m | 7 | 7 | 7 | PL | Sympatric | Collected | 50.73 | 86.34 | 17 |
| Emberiza_GBS4_XD_674 | Hybrid | f | NA | NA | NA | FML | Sympatric | Collected | 50.73 | 86.34 | 17 |
| Emberiza_GBS4_XD_684 | E. citrinella | m | 0 | 1 | 0 | SC | Sympatric | Collected | 51.57 | 85.56 | 17 |
| Emberiza_GBS4_XD_688 | Hybrid | f | NA | NA | NA | FML | Sympatric | Collected | 51.57 | 85.56 | 17 |
| Emberiza_GBS4_XD_699 | E. leucocephalos | m | 7 | 7 | 7 | PL | Sympatric | Collected | 50.64 | 87.96 | 17 |
| Emberiza_GBS4_XD_700 | E. leucocephalos | m | 7 | 7 | 7 | PL | Sympatric | Collected | 50.64 | 87.96 | 17 |
| Emberiza_GBS4_XD_792 | Hybrid | m | 4 | 2 | 6 | WH | Sympatric | Collected | 50.73 | 86.34 | 17 |
| Emberiza_GBS4_XD_798 | Hybrid | m | 0 | 0 | 5 | WH | Sympatric | Collected | 50.73 | 86.34 | 17 |
| Emberiza_GBS4_XD_800 | Hybrid | m | 3 | 0 | 5 | WH | Sympatric | Collected | 50.73 | 86.34 | 17 |
| Emberiza_GBS4_XD_930 | Hybrid | m | 7 | 7 | 1 | YH | Sympatric | Collected | 50.31 | 87.6 | 17 |
| Emberiza_GBS4_XD_934 | Hybrid | m | 5 | 5 | 5 | LH | Sympatric | Collected | 50.73 | 86.34 | 17 |
| Emberiza_GBS4_XD_940 | E. leucocephalos | m | 7 | 7 | 7 | PL | Sympatric | Collected | 50.73 | 86.34 | 17 |
| Emberiza_GBS4_XD_941 | Hybrid | m | 3 | 1 | 0 | CH | Sympatric | Collected | 50.73 | 86.34 | 17 |
| Emberiza_GBS4_XD_946 | Hybrid | m | 4 | 4 | 5 | LH | Sympatric | Collected | 50.73 | 86.34 | 17 |
| Emberiza_GBS4_XD_947 | Hybrid | m | 6 | 3 | 1 | YH | Sympatric | Collected | 50.73 | 86.34 | 17 |
| Emberiza_GBS4_XD_948 | E. citrinella | m | 0 | 0 | 0 | PC | Sympatric | Collected | 50.73 | 86.34 | 17 |
| Emberiza_GBS4_XD_949 | Hybrid | m | 0 | 0 | 5 | WH | Sympatric | Collected | 50.73 | 86.34 | 17 |
| Emberiza_GBS4_XD_950 | Hybrid | m | 5 | 3 | 1 | YH | Sympatric | Collected | 50.73 | 86.34 | 17 |
| Emberiza_GBS4_XD_957 | Hybrid | m | 4 | 3 | 0 | CH | Sympatric | Collected | 50.73 | 86.34 | 17 |
| Emberiza_GBS4_XD_958 | E. citrinella | f | NA | NA | NA | FML | Sympatric | Collected | 50.73 | 86.34 | 17 |
| Emberiza_GBS4_XD_960 | Hybrid | m | 7 | 7 | 0 | YH | Sympatric | Collected | 50.73 | 86.34 | 17 |
| Emberiza_GBS4_XD_961 | Hybrid | m | 3 | 1 | 0 | CH | Sympatric | Collected | 50.73 | 86.34 | 17 |
| Emberiza_GBS4_XD_962 | Hybrid | m | 7 | 6 | 0 | YH | Sympatric | Collected | 50.73 | 86.34 | 17 |
| Emberiza_GBS4_XD_967 | E. leucocephalos | m | 7 | 7 | 7 | PL | Sympatric | Collected | 50.73 | 86.34 | 17 |

| Sample ID | Species | Sex | TH | BR | BG | Pheno Class | Geographic Distribution | Source | Latitude (°N) | Longitude (°E) | Sampling Location |
|---|---|---|---|---|---|---|---|---|---|---|---|
| Emberiza_GBS4_XD_968 | Hybrid | m | 3 | 2 | 0 | CH | Sympatric | Collected | 50.73 | 86.34 | 17 |
| Emberiza_GBS4_XD_971 | Hybrid | m | 5 | 4 | 7 | LH | Sympatric | Collected | 50.73 | 86.34 | 17 |
| Emberiza_GBS4_XD_972 | E. leucocephalos | m | 7 | 7 | 7 | PL | Sympatric | Collected | 50.73 | 86.34 | 17 |
| Emberiza_GBS4_XD_976 | Hybrid | m | 3 | 2 | 1 | CH | Sympatric | Collected | 50.73 | 86.34 | 17 |
| Emberiza_GBS4_XD_977 | Hybrid | m | 3 | 2 | 0 | CH | Sympatric | Collected | 50.73 | 86.34 | 17 |
| Emberiza_GBS4_XD_979 | Hybrid | m | 7 | 7 | 0 | YH | Sympatric | Collected | 50.73 | 86.34 | 17 |
| Emberiza_GBS4_XD_990 | E. citrinella | m | 0 | 0 | 0 | PC | Sympatric | Collected | 50.73 | 86.34 | 17 |
| Emberiza_GBS4_XD_994 | Hybrid | m | 5 | 1 | 0 | CH | Sympatric | Collected | 50.73 | 86.34 | 17 |
| Emberiza_GBS4_XD_995 | Hybrid | m | 2 | 3 | 7 | WH | Sympatric | Collected | 50.73 | 86.34 | 17 |
| Emberiza_GBS4_XD_997 | E. citrinella | m | 2 | 1 | 0 | SC | Sympatric | Collected | 50.73 | 86.34 | 17 |
| Emberiza_GBS4_ZM_1164 | E. citrinella | m | 1 | 0 | 0 | SC | Sympatric | Zoological museum, Russia | 54.93 | 86.82 | 18 |
| Emberiza_GBS4_ZM_1184 | E. citrinella | m | 1 | 0 | 0 | SC | Sympatric | Zoological museum, Russia | 54.93 | 86.82 | 18 |
| Emberiza_GBS4_ZM_1335 | Hybrid | m | 3 | 3 | 7 | WH | Sympatric | Zoological museum, Russia | 51.88 | 80.1 | 15 |
| Emberiza_GBS5_ASR05_62_2 | E. leucocephalos | m | 7 | 7 | 7 | PL | Allopatric | Collected | 50.21 | 115.06 | 28 |
| Emberiza_GBS5_ENP05_02 | E. citrinella | uk | NA | NA | NA | UK | Sympatric | Collected | 51.96 | 85.96 | 17 |
| Emberiza_GBS5_RYA_3003 | E. leucocephalos | m | NA | NA | NA | UK | Allopatric | Zoological museum, Russia | 50.68 | 142.97 | 30 |
| Emberiza_GBS5_SVN_2336 | E. leucocephalos | f | NA | NA | NA | FML | Allopatric | Zoological museum, Russia | 50.68 | 142.97 | 30 |
| Emberiza_GBS5_XD_549 | Hybrid | m | 7 | 7 | 1 | YH | Sympatric | Collected | 50.73 | 86.34 | 17 |

| Sample ID | Species | Sex | TH | BR | BG | Pheno Class | Geographic Distribution | Source | Latitude (°N) | Longitude (°E) | Sampling Location |
|---|---|---|---|---|---|---|---|---|---|---|---|
| Emberiza_GBS5_XD_550 | Hybrid | m | 5 | 3 | 0 | YH | Sympatric | Collected | 50.73 | 86.34 | 17 |
| Emberiza_GBS5_XD_602 | E. citrinella | m | 0 | 0 | 0 | PC | Sympatric | Collected | 50.73 | 86.34 | 17 |
| Emberiza_GBS5_XD_605 | E. leucocephalos | m | 7 | 7 | 4 | SL | Sympatric | Collected | 50.73 | 86.34 | 17 |
| Emberiza_GBS5_XD_608 | Hybrid | m | 4 | 1 | 5 | WH | Sympatric | Collected | 50.73 | 86.34 | 17 |
| Emberiza_GBS5_XD_609 | E. leucocephalos | m | 7 | 7 | 6 | SL | Sympatric | Collected | 50.73 | 86.34 | 17 |
| Emberiza_GBS5_XD_611 | Hybrid | m | 3 | 3 | 0 | CH | Near Sympatric | Collected | 55.22 | 65.42 | 14 |
| Emberiza_GBS5_XD_615 | E. citrinella | m | 0 | 0 | 0 | PC | Near Sympatric | Collected | 55.22 | 65.42 | 14 |
| Emberiza_GBS5_XD_616 | E. citrinella | m | 0 | 0 | 0 | PC | Near Sympatric | Collected | 55.22 | 65.42 | 14 |
| Emberiza_GBS5_XD_620 | Hybrid | m | 3 | 4 | 0 | CH | Near Sympatric | Collected | 55.19 | 65.26 | 14 |
| Emberiza_GBS5_XD_622 | E. citrinella | m | 0 | 0 | 0 | PC | Near Sympatric | Collected | 55.19 | 65.26 | 14 |
| Emberiza_GBS5_XD_626 | E. citrinella | m | 0 | 0 | 0 | PC | Near Sympatric | Collected | 54.67 | 64.88 | 14 |
| Emberiza_GBS5_XD_627 | Hybrid | m | 6 | 6 | 0 | YH | Near Sympatric | Collected | 54.67 | 64.88 | 14 |
| Emberiza_GBS5_XD_628 | Hybrid | m | 7 | 7 | 1 | YH | Near Sympatric | Collected | 54.67 | 64.88 | 14 |
| Emberiza_GBS5_XD_629 | E. leucocephalos | m | 7 | 7 | 7 | PL | Near Sympatric | Collected | 54.67 | 64.88 | 14 |
| Emberiza_GBS5_XD_633 | Hybrid | m | 6 | 5 | 0 | YH | Sympatric | Collected | 50.73 | 86.34 | 17 |
| Emberiza_GBS5_XD_638 | Hybrid | m | 1 | 0 | 6 | WH | Sympatric | Collected | 50.73 | 86.34 | 17 |
| Emberiza_GBS5_XD_642 | Hybrid | m | 3 | 2 | 0 | CH | Sympatric | Collected | 50.73 | 86.34 | 17 |
| Emberiza_GBS5_XD_643 | Hybrid | m | 2 | 1 | 5 | WH | Sympatric | Collected | 50.73 | 86.34 | 17 |

| Sample ID | Species | Sex | TH | BR | BG | Pheno Class | Geographic Distribution | Source | Latitude (°N) | Longitude (°E) | Sampling Location |
|---|---|---|---|---|---|---|---|---|---|---|---|
| Emberiza_GBS5_XD_646 | E. leucocephalos | m | 7 | 7 | 7 | PL | Sympatric | Collected | 50.73 | 86.34 | 17 |
| Emberiza_GBS5_XD_648 | E. citrinella | m | 1 | 1 | 0 | SC | Sympatric | Collected | 50.73 | 86.34 | 17 |
| Emberiza_GBS5_XD_651 | E. leucocephalos | m | 7 | 7 | 7 | PL | Sympatric | Collected | 50.73 | 86.34 | 17 |
| Emberiza_GBS5_XD_660 | E. citrinella | m | 1 | 1 | 0 | SC | Allopatric | Collected | 56.06 | 36.13 | 7 |
| Emberiza_GBS5_XD_661 | E. citrinella | m | 0 | 0 | 0 | PC | Allopatric | Collected | 56.06 | 36.13 | 7 |
| Emberiza_GBS5_XD_667 | Hybrid | m | 4 | 4 | 1 | CH | Sympatric | Collected | 50.73 | 86.34 | 17 |
| Emberiza_GBS5_XD_668 | Hybrid | f | NA | NA | NA | FML | Sympatric | Collected | 50.73 | 86.34 | 17 |
| Emberiza_GBS5_XD_683 | E. citrinella | m | 1 | 0 | 0 | SC | Sympatric | Collected | 53.09 | 83.84 | 16 |
| Emberiza_GBS5_XD_685 | E. citrinella | m | 1 | 1 | 1 | SC | Sympatric | Collected | 51.57 | 85.56 | 17 |
| Emberiza_GBS5_XD_686 | Hybrid | m | 5 | 5 | 7 | LH | Sympatric | Collected | 51.57 | 85.56 | 17 |
| Emberiza_GBS5_XD_687 | E. citrinella | m | 1 | 0 | 0 | SC | Sympatric | Collected | 51.57 | 85.56 | 17 |
| Emberiza_GBS5_XD_689 | Hybrid | m | 4 | 3 | 0 | CH | Sympatric | Collected | 51.57 | 85.56 | 17 |
| Emberiza_GBS5_XD_690 | Hybrid | m | 1 | 1 | 5 | WH | Sympatric | Collected | 53.09 | 83.84 | 16 |
| Emberiza_GBS5_XD_694 | Hybrid | m | 6 | 5 | 7 | LH | Sympatric | Collected | 50.64 | 87.96 | 17 |
| Emberiza_GBS5_XD_698 | E. leucocephalos | m | 7 | 7 | 7 | PL | Sympatric | Collected | 50.64 | 87.96 | 17 |
| Emberiza_GBS5_XD_794 | E. leucocephalos | m | 7 | 7 | 7 | PL | Sympatric | Collected | 50.73 | 86.34 | 17 |
| Emberiza_GBS5_XD_795 | E. citrinella | m | 0 | 0 | 1 | PC | Sympatric | Collected | 50.73 | 86.34 | 17 |
| Emberiza_GBS5_XD_797 | E. leucocephalos | m | 6 | 6 | 6 | SL | Sympatric | Collected | 50.73 | 86.34 | 17 |
| Emberiza_GBS5_XD_799 | E. citrinella | m | 0 | 0 | 0 | PC | Sympatric | Collected | 50.73 | 86.34 | 17 |
| Emberiza_GBS5_XD_924 | E. leucocephalos | m | 7 | 7 | 7 | PL | Sympatric | Collected | 50.02 | 89.23 | 19 |
| Emberiza_GBS5_XD_925 | E. leucocephalos | m | 7 | 7 | 7 | PL | Sympatric | Collected | 50.02 | 89.23 | 19 |
| Emberiza_GBS5_XD_927 | E. leucocephalos | m | 7 | 7 | 6 | SL | Sympatric | Collected | 50.31 | 87.6 | 17 |
| Emberiza_GBS5_XD_928 | E. citrinella | m | 2 | 1 | 0 | SC | Sympatric | Collected | 50.31 | 87.6 | 17 |
| Emberiza_GBS5_XD_931 | Hybrid | m | 4 | 1 | 0 | CH | Sympatric | Collected | 50.73 | 86.34 | 17 |

| Sample ID | Species | Sex | TH | BR | BG | Pheno Class | Geographic Distribution | Source | Latitude (°N) | Longitude (°E) | Sampling Location |
|---|---|---|---|---|---|---|---|---|---|---|---|
| Emberiza_GBS5_XD_942 | E. citrinella | m | 1 | 0 | 0 | SC | Sympatric | Collected | 50.73 | 86.34 | 17 |
| Emberiza_GBS5_XD_943 | E. citrinella | m | 1 | 1 | 0 | SC | Sympatric | Collected | 50.73 | 86.34 | 17 |
| Emberiza_GBS5_XD_944 | Hybrid | m | 3 | 1 | 0 | CH | Sympatric | Collected | 50.73 | 86.34 | 17 |
| Emberiza_GBS5_XD_952 | E. leucocephalos | m | 7 | 7 | 7 | PL | Sympatric | Collected | 50.73 | 86.34 | 17 |
| Emberiza_GBS5_XD_954 | E. citrinella | m | 0 | 0 | 0 | PC | Sympatric | Collected | 50.73 | 86.34 | 17 |
| Emberiza_GBS5_XD_955 | Hybrid | m | 5 | 3 | 7 | LH | Sympatric | Collected | 50.73 | 86.34 | 17 |
| Emberiza_GBS5_XD_956 | E. citrinella | m | 2 | 1 | 0 | SC | Sympatric | Collected | 50.73 | 86.34 | 17 |
| Emberiza_GBS5_XD_963 | Hybrid | m | 0 | 0 | 7 | WH | Sympatric | Collected | 50.73 | 86.34 | 17 |
| Emberiza_GBS5_XD_964 | E. leucocephalos | m | 7 | 7 | 5 | SL | Sympatric | Collected | 50.73 | 86.34 | 17 |
| Emberiza_GBS5_XD_965 | E. citrinella | f | NA | NA | NA | FML | Sympatric | Collected | 50.73 | 86.34 | 17 |
| Emberiza_GBS5_XD_966 | E. citrinella | m | 2 | 1 | 0 | SC | Sympatric | Collected | 50.73 | 86.34 | 17 |
| Emberiza_GBS5_XD_969 | Hybrid | m | 5 | 5 | 5 | LH | Sympatric | Collected | 50.73 | 86.34 | 17 |
| Emberiza_GBS5_XD_970 | E. leucocephalos | m | 7 | 7 | 7 | PL | Sympatric | Collected | 50.73 | 86.34 | 17 |
| Emberiza_GBS5_XD_973 | E. citrinella | m | 0 | 0 | 0 | PC | Sympatric | Collected | 50.73 | 86.34 | 17 |
| Emberiza_GBS5_XD_974 | Hybrid | m | 1 | 1 | 5 | WH | Sympatric | Collected | 50.73 | 86.34 | 17 |
| Emberiza_GBS5_XD_975 | E. leucocephalos | m | 7 | 7 | 7 | PL | Sympatric | Collected | 50.73 | 86.34 | 17 |
| Emberiza_GBS5_XD_978 | E. citrinella | m | 1 | 1 | 0 | SC | Sympatric | Collected | 50.73 | 86.34 | 17 |
| Emberiza_GBS5_XD_980 | E. leucocephalos | m | 6 | 7 | 4 | SL | Sympatric | Collected | 50.73 | 86.34 | 17 |
| Emberiza_GBS5_XD_981 | E. leucocephalos | f | NA | NA | NA | FML | Sympatric | Collected | 50.73 | 86.34 | 17 |
| Emberiza_GBS5_XD_982 | E. citrinella | m | 0 | 0 | 1 | PC | Sympatric | Collected | 50.73 | 86.34 | 17 |
| Emberiza_GBS5_XD_983 | Hybrid | m | 3 | 1 | 0 | CH | Sympatric | Collected | 50.73 | 86.34 | 17 |
| Emberiza_GBS5_XD_984 | E. citrinella | m | 2 | 1 | 0 | SC | Sympatric | Collected | 50.73 | 86.34 | 17 |
| Emberiza_GBS5_XD_985 | E. leucocephalos | m | 6 | 6 | 7 | SL | Sympatric | Collected | 50.73 | 86.34 | 17 |
| Emberiza_GBS5_XD_987 | E. citrinella | m | 0 | 0 | 0 | PC | Sympatric | Collected | 50.73 | 86.34 | 17 |

| Sample ID | Species | Sex | TH | BR | BG | Pheno Class | Geographic Distribution | Source | Latitude (°N) | Longitude (°E) | Sampling Location |
|---|---|---|---|---|---|---|---|---|---|---|---|
| Emberiza_GBS5_XD_989 | E. leucocephalos | m | 7 | 7 | 5 | SL | Sympatric | Collected | 50.73 | 86.34 | 17 |
| Emberiza_GBS5_XD_992 | E. citrinella | m | 0 | 0 | 0 | PC | Sympatric | Collected | 50.73 | 86.34 | 17 |
| Emberiza_GBS5_XD_993 | Hybrid | m | 4 | 3 | 1 | CH | Sympatric | Collected | 50.73 | 86.34 | 17 |
| Emberiza_GBS5_XD_999 | Hybrid | m | 3 | 0 | 1 | CH | Sympatric | Collected | 50.73 | 86.34 | 17 |
| Emberiza_GBS5_ZM_1162 | E. citrinella | m | 1 | 0 | 0 | PC | Sympatric | Zoological museum, Russia | 54.93 | 86.82 | 18 |
| Emberiza_GBS5_ZM_1183 | Hybrid | m | 7 | 7 | 0 | YH | Sympatric | Zoological museum, Russia | 54.93 | 86.82 | 18 |
| Emberiza_GBS5_ZM_1197 | E. leucocephalos | m | 7 | 7 | 7 | PL | Sympatric | Zoological museum, Russia | 53.43 | 83.93 | 16 |
| Emberiza_GBS5_ZM_1200 | Hybrid | m | 2 | 0 | 6 | WH | Sympatric | Zoological museum, Russia | 53.43 | 83.93 | 16 |
| Emberiza_GBS5_ZM_1220 | Hybrid | m | 3 | 3 | 0 | CH | Sympatric | Zoological museum, Russia | 53.43 | 83.93 | 16 |
| Emberiza_GBS5_ZM_1267 | E. leucocephalos | m | 6 | 6 | 7 | SL | Sympatric | Zoological museum, Russia | 53.37 | 78.02 | 15 |
| Emberiza_GBS5_ZM_1375 | E. citrinella | m | 1 | 0 | 0 | PC | Sympatric | Zoological museum, Russia | 53.24 | 83.51 | 16 |

**Supplementary Table 2.** Detailed information on the three pairs of sympatric individuals that showed appreciably large kinship coefficients. Explanations for the abbreviations used in the "Pheno Class" (Phenotypic Class) column can be found in the Methods section. In the "sex" column, "m" stands for male, "f" stands for female and "uk" stands for unknown. The "TH" column contains phenotypic scores for each individual for the throat plumage trait. The "BR" column contains phenotypic scores for each individual for the brow plumage trait. The "BG" column contains phenotypic scores for each individual for the background colour plumage trait. In the "Pheno Class" column, "FML" stands for female and "UK" stands for unknown. The numbers in the "Sampling Location" column correspond to those that appear in Figure 1A. In all columns, a "NA" observation stands for "Not Applicable".

| Sample ID | Species | Sex | TH | BR | BG | Pheno. Class | Latitude (°N) | Longitude (°E) | Sampling Location | Kinship Coefficient | Kinship Relationship |
|---|---|---|---|---|---|---|---|---|---|---|---|
| **Pair 1** | | | | | | | | | | | |
| Emberiza_GBS1_VGR_508 | E. citrinella | m | 0 | 0 | 1 | SC | 56.405 | 105.463 | 25 | 0.048301968 | 3rd degree |
| Emberiza_GBS1_AWH_169 | E. citrinella | f | NA | NA | NA | FML | 56.405 | 105.463 | 25 | | |
| **Pair 2** | | | | | | | | | | | |
| Emberiza_GBS1_ENP05_06 | E. citrinella | uk | NA | NA | NA | UK | 51.96 | 85.96 | 17 | 0.2323328 | 1st degree |
| Emberiza_GBS5_XD_689 | Hybrid | m | 4 | 3 | 0 | CH | 51.57 | 85.56 | 17 | | |
| **Pair 3** | | | | | | | | | | | |
| Emberiza_GBS2_ENP05_12 | E. leucocephalos | uk | NA | NA | NA | UK | 51.96 | 85.96 | 17 | 0.3095717 | 1st degree |
| Emberiza_GBS5_ENP05_02 | E. citrinella | uk | NA | NA | NA | UK | 51.96 | 85.96 | 17 | | |
